# Axon development is regulated at genetic and proteomic interfaces between the integrin adhesome and the RPM-1 ubiquitin ligase signaling hub

**DOI:** 10.1101/2023.11.15.566604

**Authors:** Jonathan Amezquita, Muriel Desbois, Karla J. Opperman, Joseph S. Pak, Elyse L. Christensen, Nikki T. Nguyen, Karen Diaz-Garcia, Melissa A. Borgen, Brock Grill

**Author notes:** These authors contributed equally (Jonathan Amezquita, Muriel Desbois).

## Abstract

Integrin signaling plays important roles in development and disease. An adhesion signaling network called the integrin adhesome has been principally defined using bioinformatics and proteomics. To date, the adhesome has not been studied using integrated proteomic and genetic approaches. Here, proteomic studies in *C. elegans* identified physical associations between the RPM-1 ubiquitin ligase signaling hub and numerous adhesome components including Talin, Kindlin and beta-integrin. *C. elegans* RPM-1 is orthologous to human MYCBP2, a prominent player in nervous system development associated with a neurodevelopmental disorder. Using neuron-specific, CRISPR loss-of-function strategies, we show that core adhesome components affect axon development and interact genetically with RPM-1. Mechanistically, Talin opposes RPM-1 in a functional ‘tug-of-war’ on growth cones that is required for accurate axon termination. Thus, our findings orthogonally validate the adhesome via multi-component genetic and physical interfaces with a key neuronal signaling hub and identify new links between the adhesome and brain disorders.

## INTRODUCTION

In the nervous system, integrin receptors and their signaling mechanisms have broad functions in cell migration, axon development, synaptic connectivity and axon regeneration ^1–4^. Integrin signaling has been linked to neurological conditions such as autism spectrum disorder (ASD) and Alzheimer’s disease ^2, 3^. Outside the nervous system, integrin signaling plays prominent roles in tissue development, the immune system, and cancer ^5–7^.

Integrins are transmembrane receptors in a large signaling and adhesion network referred to as the integrin adhesome ^8–11^. The integrin adhesome facilitates cellular interactions with the extracellular matrix by linking ECM components to intracellular F-actin thereby influencing cell migration and cellular process outgrowth. In the nervous system, integrin signaling figures prominently in neurite and axon outgrowth and guidance ^12–16^. Talin and Kindlin are two core components of the adhesome which bind beta-integrin receptors ^17, 18^. Prior studies have examined Talin and Kindlin in axon development using cultured neurons and by targeting upstream regulatory players such as calpain ^19, 20^. To date, studies that genetically impair Talin or Kindlin directly and examine effects on axon and growth cone development in vivo remain absent. Moreover, the identification of the integrin adhesome is primarily based on bioinformatics and proteomics ^8^. We are unaware of studies that have integrated proteomics and genetics to evaluate how multiple integrin adhesome components influence neuronal development in an organismal setting. An invertebrate genetic model organism, such as the nematode *C. elegans*, is ideal for such studies because of its genetic tractability and simplified integrin signaling with a single gene encoding β-integrin (PAT-3), Talin (TLN-1) and Kindlin (UNC-112).

While it is essential for axons to grow and be successfully guided to target cells, it is also critical that outgrowth be terminated – a process called axon termination. Efficient, precise nervous system construction requires spatially and temporally accurate axon termination. For example, proper axon termination is required to generate axonal tiling patterns that occur in laminated termination zones in the spinal cord, cortical layers and retina ^21^. The importance of understanding axon termination is further highlighted by ongoing efforts to comprehensively map axon termination patterns in the rodent brain ^22^. At present, it remains unclear whether adhesome signaling influences axon termination.

One of the most prominent players in axon termination identified using *C. elegans* is the Regulator of Presynaptic Morphology-1 (RPM-1) ^23–26^. *C. elegans* RPM-1 is a ubiquitin ligase signaling hub that is orthologous to human MYCBP2 ^23, 27–29^. MYCBP2 (also called Phr1 in rodents) is an evolutionarily conserved regulator of axon development ^30, 31^, and genetic variants in *MYCBP2* were recently shown to cause a neurodevelopmental spectrum disorder called *MYCBP2*-related developmental delay with corpus callosum defects (MDCD) ^32^. Despite the importance of RPM-1/MYCBP2 in nervous system development and disease, we know very little about genetic factors that oppose RPM-1 during growth cone and axon development. Moreover, physical and genetic interactions between RPM-1 and the integrin adhesome have not been examined in any system to our knowledge.

Using unbiased, affinity purification (AP) proteomics with RPM-1, we identified numerous conserved components of the *C. elegans* integrin adhesome physically associated with RPM-1. Interestingly, we found genetic links to neurobehavioral abnormalities for 75% of the components in the adhesome subnetwork we identified. CRISPR-based, cell-specific loss of function strategies demonstrated that impairing core adhesome components, PAT-3/β-integrin, TLN-1/Talin and UNC-112/Kindlin, resulted in premature axon termination in mechanosensory neurons. Genetic interaction studies and developmental time-course results indicate that the adhesome engages in a functional ‘tug-of-war’ with RPM-1 during growth cone development. Developing animals require this genetic interface between the adhesome and RPM-1 to yield anatomically accurate and temporally precise axon termination.

## RESULTS

### AP-proteomics identifies physical associations between numerous integrin adhesome components and RPM-1

To identify proteins physically associated with the RPM-1 ubiquitin ligase signaling hub, we previously performed large-scale, unbiased AP-proteomics using *C. elegans*. We relied upon integrated transgenes that express RPM-1 fused with a Protein G::Streptavidin binding peptide (GS) affinity tag expressed using the native *rpm-1* promoter on an *rpm-1* protein null background ^25, 33^. Negative control animals expressed an integrated transgene where the GS tag was fused to GFP (GS::GFP) and expressed by the *rpm-1* promoter. We previously established two groups of RPM-1 associated proteins identified by AP-proteomics, RPM-1 binding proteins and RPM-1 ubiquitination substrates (**Fig 1A**) ^25, 34^. We previously showed that RPM-1 binding proteins, such as RAE-1, GLO-4, FSN-1 and PPM-2, are present in both the GS::RPM-1 and GS::RPM-1 LD samples compared to the GS::GFP negative control **(Fig 1B**) ^25, 34–37^. In contrast, the RPM-1 LD biochemically ‘traps’ and enriches ubiquitination substrates, such as ULK/UNC-51 and CDK-5 (**Fig 1C**) ^25, 34^.

**Figure 1.**
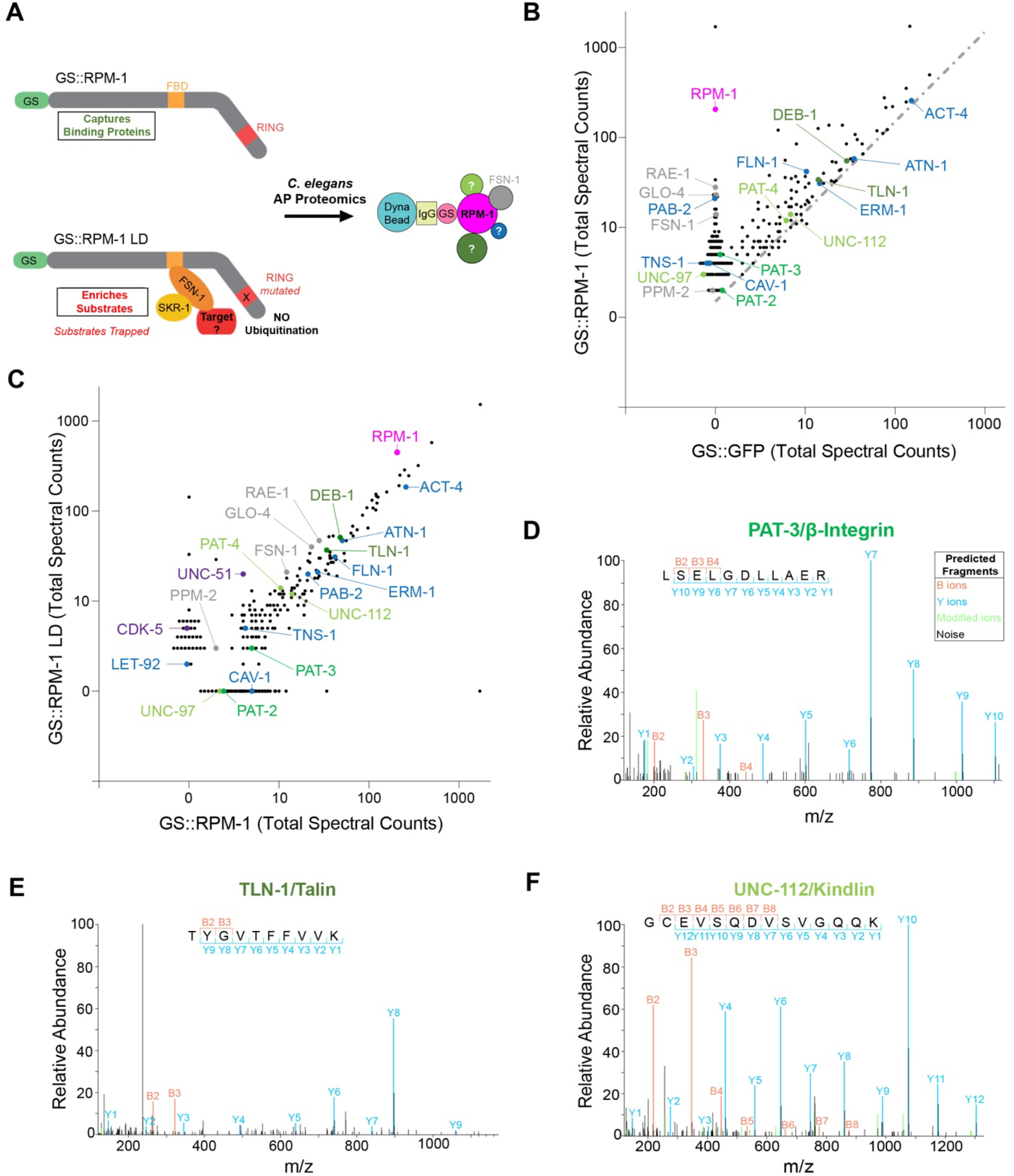
RPM-1 AP-proteomics identifies several integrin adhesome components. **A)** Schematic of RPM-1 ubiquitin ligase constructs used for AP-proteomics from *C. elegans*. Both GS::RPM-1 and GS::RPM-1 LD capture binding proteins (e.g. FSN-1), while GS::RPM-1 LD enriches ubiquitination substrates. **B)** Example of single AP-proteomics experiment showing individual proteins identified by LC-MS/MS. Shown are results from GS::RPM-1 sample compared with GS::GFP (negative control). Highlighted (shades of green) are integrin receptors, PAT-3/β-Integrin and PAT-2/α-Integrin, and two canonical integrin signaling complexes, UNC-112/Kindlin complex and TLN-1/Talin complex. Additional adhesome components (blue) were identified. Also highlighted are RPM-1 (magenta) and known RPM-1 binding proteins (gray). Dashed line delineates 1.5x fold enrichment. **C)** Results from single AP-proteomics experiment showing GS::RPM-1 compared to GS::RPM-1 LD ubiquitination substrate ‘trap’. Highlighted are previously validated substrates (CDK-5 and UNC-51, purple). **D-F)** Examples of LC-MS/MS peptide spectrum for **D)** PAT-3/β-Integrin, **E)** TLN-1/Talin, and **F)** UNC-112/Kindlin.

Here, we further analyzed our large-scale proteomics dataset, and identified numerous components of the integrin adhesome physically associated with RPM-1 (**Fig 1B-F**). While a prior bioinformatic study demonstrated that the adhesome is conserved in *C. elegans* ^38^, we curated and updated the *C. elegans* adhesome based on several criteria: 30% or greater protein sequence similarity to mammalian adhesome components, 20% or greater protein sequence identity, and conserved protein domains. Our analysis identified 132 mammalian adhesome components in *C. elegans* with 123 components having unique nematode orthologs (**Table S1**).

In total, we identified 32 *C. elegans* integrin adhesome components enriched more than 1.5 times in GS::RPM-1 samples compared to GS::GFP negative control samples (**Fig 1B, D-F; Table 1**). Adhesome components were not enriched in the GS::RPM-1 LD sample compared to the GS::RPM-1 sample (**Fig 1C**, **Table 1**). These results suggest that the majority of adhesome components associated with RPM-1 are not likely to be ubiquitination substrates. Further analysis revealed several core adhesome components and complexes associated with RPM-1. This included 1) the β-integrin PAT-3 and the α-integrin PAT-2; 2) the Kindlin complex composed of UNC-112/Kindlin, PAT-4/ILK, PAT-6/PARVA and UNC-97/LIMS, and 3) the Talin complex that contains TLN-1/Talin and DEB-1/Vinculin (**Fig 1B-F**; **Table 1**; **Fig 2A**). Several further adhesome components were also identified (**Fig 1B, C; Table 1**). Analysis of results from 7 independent proteomic experiments showed that the majority of adhesome components were identified in two or more experiments with several significantly enriched in GS::RPM-1 or GS::RPM-1 LD samples compared to GS::GFP negative controls (**Table 1**). Interestingly, significantly enriched core adhesome components included the sole *C. elegans* β-integrin PAT-3 (13% sequence coverage, **Fig S2**), the sole Talin TLN-1 (28% sequence coverage, **Fig S3**), and the sole Kindlin UNC-112 (35% sequence coverage, **Fig S4**). Thus, in vivo AP-proteomics using *C. elegans* identified physical associations between RPM-1 and numerous adhesome components.

**Figure 2.**
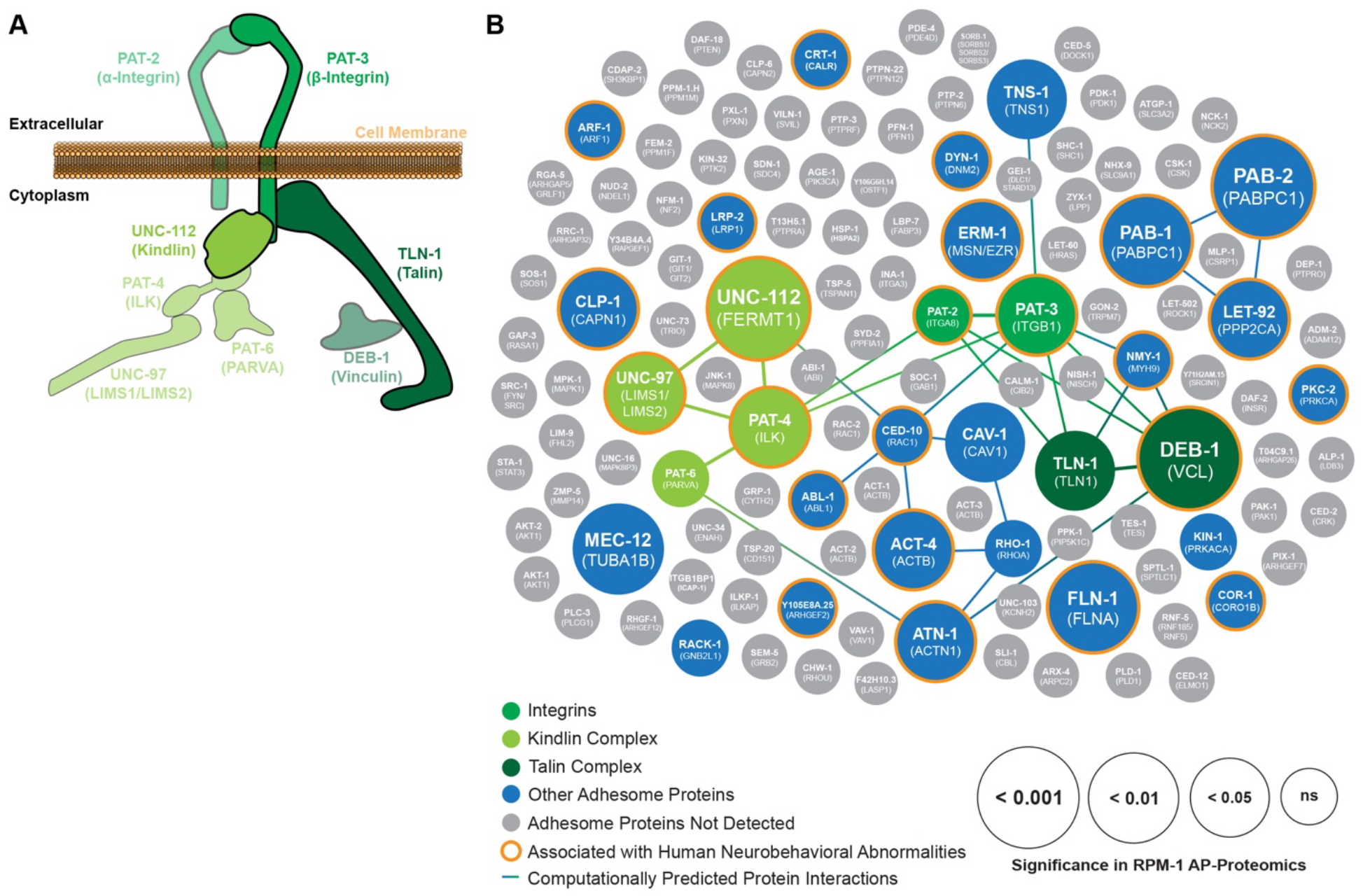
Two core adhesome signaling complexes and numerous adhesome components with links to human neurobehavioral deficits are present in RPM-1 proteomics. **A)** Illustration of integrin signaling with the two transmembrane receptors PAT-3 β-Integrin and PAT-2 α-Integrin (green), the TLN-1 Talin complex (dark green) and the UNC-112 Kindlin complex (light green). **B)** Summary of RPM-1 AP-proteomics data from *C. elegans* integrated with a computationally predicted protein-protein interaction network (lines) and genetic links to human neurobehavioral abnormalities (orange halo). Adhesome components enriched in RPM-1 AP-proteomics are highlighted in color with increasing circle size denoting significance (GS::RPM-1 test samples versus GS::GFP negative controls). Highlighted (shades of green) are Integrins, UNC-112 Kindlin complex and TLN-1 Talin complex. Also shown are additional adhesome components detected (blue) or absent (gray) in RPM-1 AP-proteomics. Orthologous human protein annotated in brackets. Data is presented from 7 independent RPM-1 AP-proteomics experiments. Significance determined using Student’s *t-test*. ns = not significant

**Table 1.** Summary of adhesome components detected in RPM-1 AP-proteomics using *C. elegans*. Shown are cumulative results from seven independent RPM-1 AP-proteomics experiments. Reported are total peptide spectra for adhesome components enriched 1.5x in either GS::RPM-1 or GS::RPM-1 LD over GS::GFP negative control samples. Note adhesome components do not show more than 2 fold increase in GS::RPM-1 LD compared to GS::RPM-1 samples suggesting they are not likely to be ubiquitination substrates. Significance determined using Student’s *t-*test. For statistical comparison and fold enrichment, normalized values (see methods) are used. ***p<0.001, **p<0.01, ns = not significant, and na = not applicable due to detection in single proteomics experiment.

|  | Protein Identified LC-MS/MS | Vertebrate Homolog | MW (kDa) | Total Experiments Enriched | Total Peptide Spectra |  |  | Significance vs GS::GFP (p-value) | RPM-1 LD vs RPM-1 Fold Increase |
| --- | --- | --- | --- | --- | --- | --- | --- | --- | --- |
|  |  |  |  |  | GS::GFP Control | GS::RPM-1 | GS::RPM-1 LD |  |  |
| Purification Target | RPM-1 | Pam/MYCBP2 | 418 | 7 | 0 | 1265 | 2874 | - | - |
| Integrin Receptors | PAT-3 | ITGB | 90 | 5 | 0 | 18 | 11 | * | 0.23x |
|  | PAT-2 | ITGA | 136 | 2 | 0 | 7 | 0 | ns | 0 |
| Kindlin Complex | UNC-112 | FERMT1 | 82 | 5 | 9 | 33 | 29 | *** | 0.32x |
|  | UNC-97 | LIMS1/LIMS2 | 40 | 3 | 0 | 9 | 0 | * | 0 |
|  | PAT-4 | ILK | 52 | 4 | 18 | 46 | 36 | * | 0.25x |
|  | PAT-6 | PARVA | 43 | 3 | 4 | 15 | 14 | ns | 0.36x |
| Talin Complex | TLN-1 | TLN1 | 279 | 5 | 36 | 101 | 113 | * | 0.47x |
|  | DEB-1 | VCL | 121 | 7 | 124 | 243 | 232 | *** | 0.35x |
| Other Adhesome Proteins | PAB-2 | PABPC1 | 76 | 6 | 0 | 128 | 143 | *** | 0.49x |
|  | FLN-1 | FLNA | 246 | 7 | 44 | 263 | 111 | ** | 0.1x |
|  | PAB-1 | PABPC1 | 72 | 6 | 176 | 401 | 389 | ** | 0.35x |
|  | MEC-12 | TUBA1B | 50 | 2 | 0 | 30 | 0 | ** | 0 |
|  | ERM-1 | MSN/EZR | 66 | 6 | 41 | 90 | 72 | * | 0.24x |
|  | ATN-1 | ACTN1 | 107 | 5 | 81 | 151 | 83 | * | 0.07x |
|  | ACT-4 | ACTB | 42 | 5 | 626 | 1408 | 879 | * | 0.12x |
|  | LET-92 | PPP2CA | 36 | 5 | 0 | 14 | 9 | * | 0.28x |
|  | CLP-1 | CAPN2 | 84 | 4 | 6 | 21 | 17 | * | 0.26x |
|  | CAV-1 | CAV1 | 26 | 3 | 0 | 10 | 0 | * | 0 |
|  | TNS-1 | TNS1 | 147 | 3 | 0 | 14 | 11 | * | 0.3x |
|  | CRT-1 | CALR | 46 | 5 | 14 | 54 | 37 | ns | 0.18x |
|  | RHO-1 | RHOA | 22 | 4 | 4 | 20 | 8 | ns | 0.13x |
|  | KIN-1 | PRKACA | 46 | 3 | 4 | 12 | 4 | ns | 0 |
|  | RACK-1 | GNB2L1 | 36 | 3 | 115 | 245 | 190 | ns | 0.16x |
|  | ARF-1 | ARF1 | 21 | 2 | 2 | 7 | 0 | ns | 0 |
|  | DYN-1 | DNM2 | 94 | 2 | 0 | 4 | 4 | ns | 0.19x |
|  | LRP-2 | LRP1 | 540 | 2 | 0 | 0 | 17 | ns | 1.92x |
|  | NMY-1 | MYH9 | 229 | 2 | 6 | 15 | 8 | ns | 0.12x |
|  | ABL-1 | ABL1 | 138 | 1 | 3 | 0 | 5 | na | na |
|  | CED-10 | RAC1 | 21 | 1 | 0 | 3 | 0 | na | na |
|  | COR-1 | CORO1B | 67 | 1 | 0 | 0 | 3 | na | na |
|  | PKC-2 | PRKCA | 82 | 1 | 0 | 2 | 0 | na | na |
|  | Y105E8A.25 | ARHGEF2 | 179 | 1 | 0 | 3 | 0 | na | na |

### Proteomic network analysis reveals that several adhesome components associated with RPM-1 are genetically linked to human neurobehavioral deficits

Next, we sought to integrate our RPM-1 proteomics data with bioinformatic network analysis of the adhesome. To do so, we generated a computationally predicted network of adhesome components identified in RPM-1 proteomics (**Fig 2B, Table S2**). We then layered statistical significance values from RPM-1 AP-proteomic results onto the subset of adhesome components associated with RPM-1 (**Fig 2B, Table S2**). Our analysis comprehensively illustrates numerous adhesome components associated with RPM-1, how significant a hit they were in AP-proteomics, and the predicted protein-protein interaction network among these players. Our results highlight the integrin receptor cluster formed by β-integrin PAT-3/ITGB1, α-integrin PAT-2/ITGA8, the Kindlin complex cluster (UNC-112/FERMT1, PAT-4/ILK, UNC-97/LIMS, PAT-6/PARVA), and the Talin complex cluster (TLN-1/TLN1, DEB-1/VCL) (**Fig 2A, B**). Importantly, we also identified many adhesome components associated with RPM-1 that do not have predicted protein-protein interactions in *C. elegans*. These included the Filamin FLN-1/FLNA, the Ezrin Moesin ortholog ERM-1/MSN, the Calpain protease CLP-1/CAPN1, the polyadenylate-binding proteins PAB-1/PABPC1 and PAB-2/PABPC1, as well as several others (**Fig 2B**).

A recent clinical genetic study showed that mutations in *MYCBP2*, the human ortholog of RPM-1, cause a neurodevelopmental spectrum disorder called MDCD ^32^. Patients with MDCD feature variable presentation of neurobehavioral abnormalities including developmental delay (DD), intellectual disability (ID), autism spectrum disorder (ASD) and epilepsy. We, therefore, investigated if any of the adhesome components present in our RPM-1 AP-proteomics dataset were also associated with human neurobehavioral abnormalities. By surveying the published literature, we made the striking observation that 75% (24/32) of adhesome components identified in RPM-1 proteomics had human genetic variants linked to neurobehavioral abnormalities (**Fig 2B, Table S3**). This included the PAT-3 β-integrin, UNC-112/Kindlin and other players in the UNC-112 complex cluster (**Fig 2B and Table S3**). These results highlight that RPM-1 is physically associated with an enriched group of adhesome components with genetic links to human neurodevelopmental deficits. Our findings are particularly intriguing, given the causal genetic link between MYCBP2/RPM-1 and a neurodevelopmental disorder.

### CRISPR labeling demonstrates native adhesome components are expressed in mechanosensory neurons and localized to axons

Because RPM-1 is associated with a cadre of adhesome components, we wanted to determine if specific core adhesome components are expressed in neurons where RPM-1 regulates axon development. RPM-1 is well known to regulate axon termination by functioning cell-autonomously in the mechanosensory neurons of *C. elegans* ^24, 39, 40^. Therefore, we investigated if PAT-3 β-integrin, UNC-112/Kindlin and TLN-1/Talin are expressed in mechanosensory neurons. To do so, we used CRISPR engineering to introduce a GFP tag at the C-terminus of the PAT-3 β-integrin and UNC-112 (**Fig S4**). For TLN-1, we evaluated a previously generated CRISPR strain where TLN-1 was N-terminally tagged with GFP (**Fig S4**) ^41^. Previous studies demonstrated that null mutations in *pat-3* and *unc-112* are lethal due to the abnormal muscle formation ^42^. It remains unclear if a genetic null of *tln-1* is viable because *tln-1* alleles described do not eliminate all TLN-1 isoforms (**Fig S4**). Thus, viability of animals with CRISPR GFP tags on PAT-3 and UNC-112 suggests that the addition of GFP does not impair gene function. We also used AlphaFold to predict the structure of CRISPR engineered PAT-3::GFP (**Fig 3A**), UNC-112::GFP (**Fig 3B**), and GFP::TLN-1 (**Fig 3C**). In all cases, the GFP tag was not predicted to interfere with protein folding (**Fig 3A-C**). Thus, structural predictions and viability of GFP knock in strains indicate that CRISPR engineered GFP constructs are suitable for evaluating expression and localization of core adhesome components.

**Figure 3.**
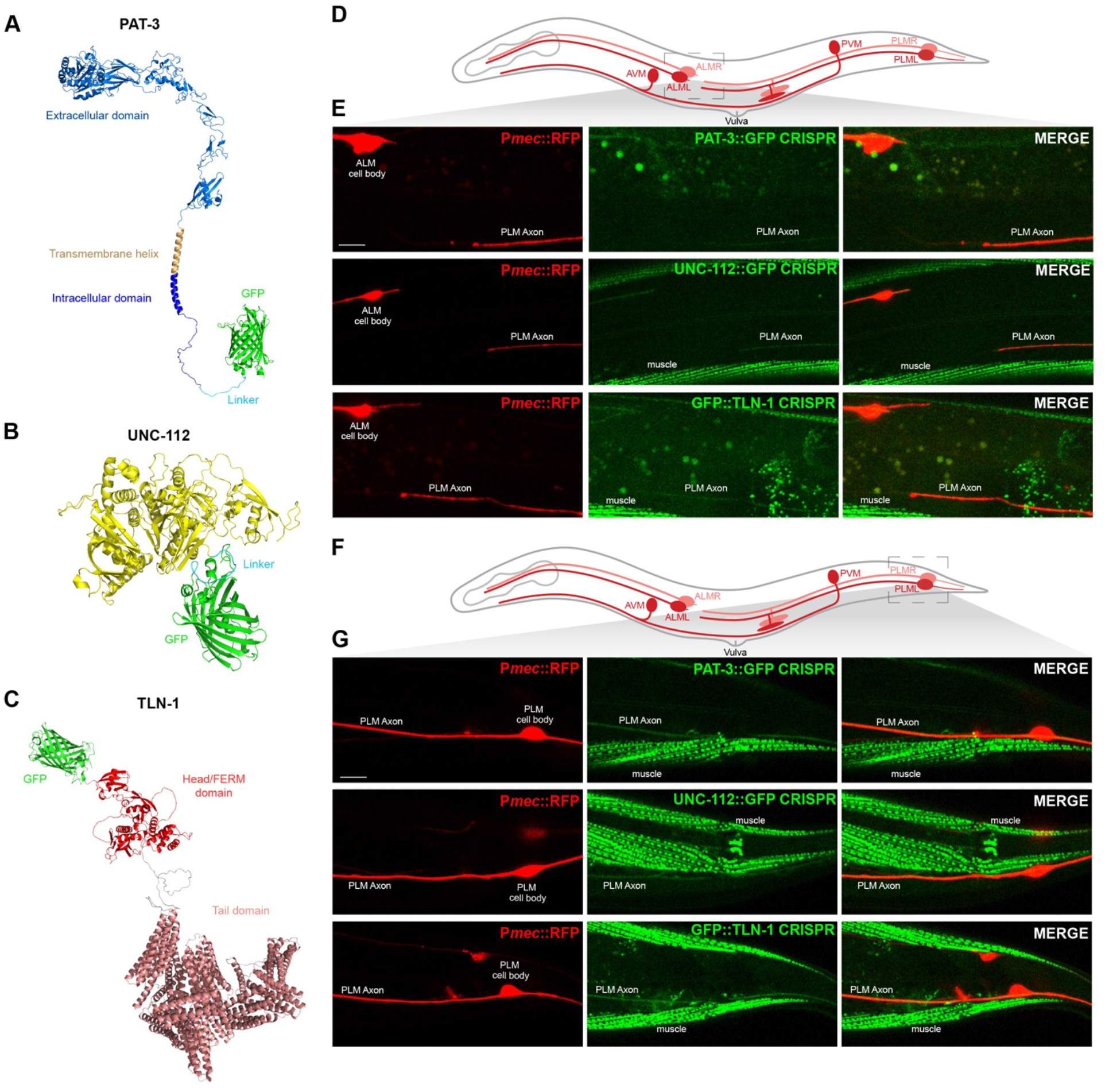
Adhesome components are expressed in *C. elegans* mechanosensory neurons and localized to axons. **A-C)** Alpha-fold predictions showing adhesome signaling components that were GFP-tagged using CRISPR engineering in *C. elegans*. Structural predictions indicate GFP is not likely to interfere with protein folding. **A)** *C. elegans* PAT-3::GFP, **B)** UNC-112::GFP, and **C)** GFP::TLN-1. **D)** Schematic of *C. elegans* mechanosensory neurons highlighting region imaged to visualize axon termination sites for PLM neurons (light gray box). **E)** Representative images showing that PAT-3::GFP, UNC-112::GFP and GFP::TLN-1 are expressed in PLM neurons and localized to axons. P*mec-7*::RFP (*jsIs973*) labels mechanosensory neurons. **F)** Schematic highlights cell body and initial axon segment (light gray box) of PLM neurons. **G)** Confocal images show PAT-3::GFP, UNC-112::GFP and GFP::TLN-1 primarily localized to PLM axon. For E and G, note that integrin components also show prominent expression in muscles. Scale bars 10μm.

*C. elegans* has two anterior ALM and two posterior PLM mechanosensory neurons located on the right and left side of its body (**Fig 3D**). Individual AVM and PVM mechanosensory neurons are on one side of the animal. Each ALM neuron has a soma in the midbody and sends a single axon toward the nose (**Fig 3D**). Each PLM soma is in the tail and has a single axon that terminates just prior to the ALM cell body (**Fig 3D**). Previous studies have shown that RPM-1 is expressed in both ALM and PLM neurons, and localized to axons and the soma ^25, 34, 39^.

We observed PAT-3::GFP, UNC-112::GFP and GFP::TLN-1 CRISPR primarily in the axons of ALM and PLM neurons (**Fig 3D-G**). Further evaluation of PLM mechanosensory neurons showed that PAT-3::GFP, UNC-112::GFP and GFP::TLN-1 are localized along the length of the axon and detected at lower levels in PLM soma (**Fig 3D-G; Fig S5**). Consistent with prior studies, we also observed PAT-3::GFP, UNC-112::GFP and GFP::TLN-1 in muscles (**Fig 3E-G**). These findings with endogenous CRISPR tagged proteins show that three core adhesome components are expressed in mechanosensory neurons, similar to RPM-1.

### CRISPR-based cell-specific degradation of adhesome components results in premature axon termination in mechanosensory neurons

Having shown that core adhesome components are expressed in mechanosensory neurons, we wanted to determine if genetic perturbation of adhesome components influences axon development. However, null mutants for *pat-3* and *unc-112* are lethal and lead to embryonic arrest at the 2-fold stage ^42^. As a result, prior genetic studies in *C. elegans* have often relied upon hypomorphic mutants. To circumvent these issues, we turned to a CRISPR-based, cell-specific approach to impair adhesome components (**Fig 4A**). We combined an anti-GFP protein degradation module expressed specifically in mechanosensory neurons (*mecDEG*) ^43^ with PAT-3::GFP, UNC-112::GFP and GFP::TLN-1 CRISPR knock in strains. This *mecDEG* system transgenically expresses the SOCS-box adaptor protein ZIF-1 fused with an anti-GFP nanobody. Anti-GFP nanobodies recognize CRISPR GFP tagged adhesome components and target them for proteasome-mediated degradation by the endogenous Rbx1/Cul2 E3 ubiquitin ligase complex. This approach allowed us to circumvent lethality and evaluate how several core adhesome components affect axon development in mechanosensory neurons.

**Figure 4.**
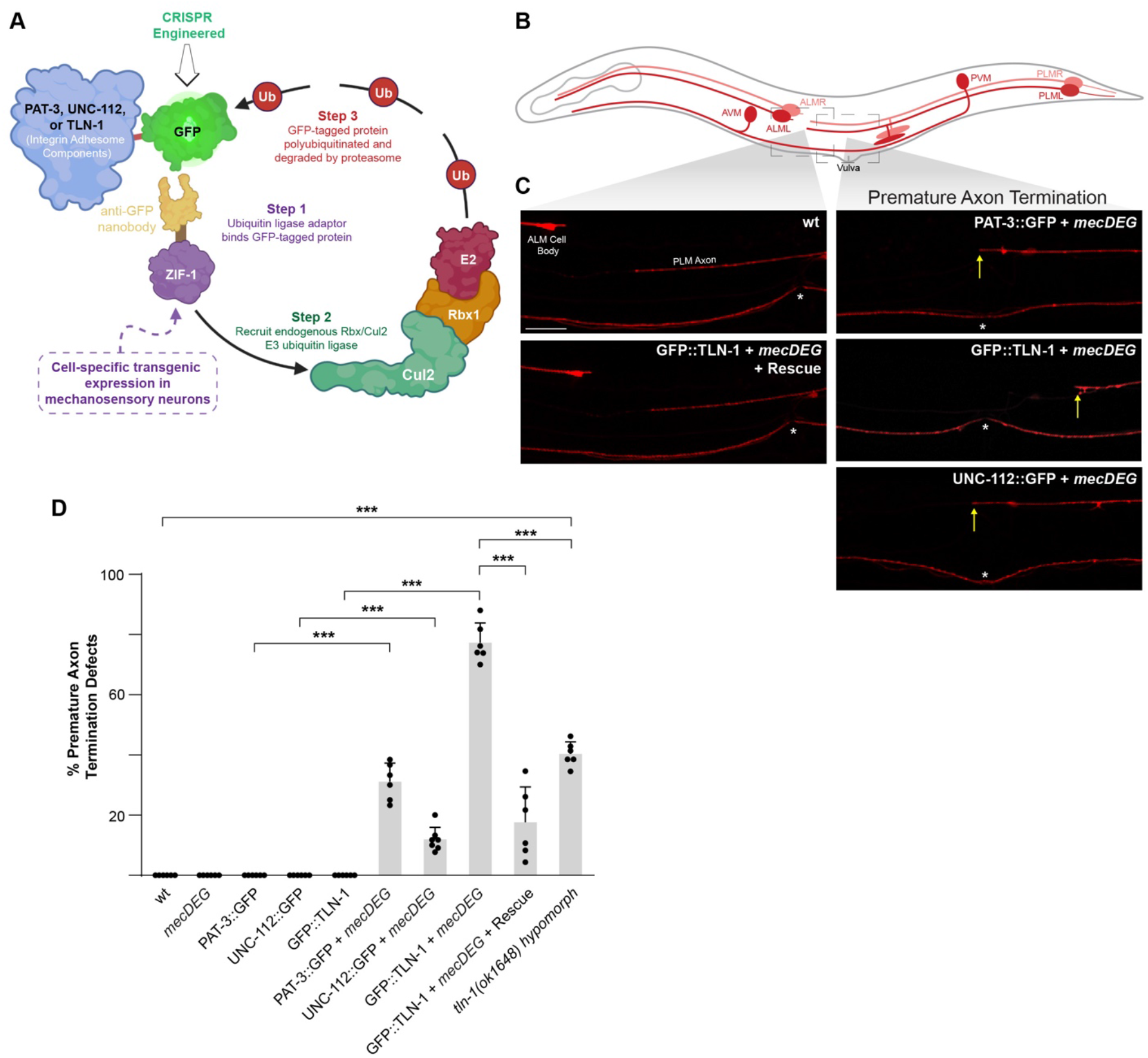
Impairing adhesome components with a CRISPR-based, cell-specific protein degradation system results in premature axon termination. **A)** Illustration of CRISPR-based cell-specific protein degradation system used to impair integrin adhesome components in *C. elegans* mechanosensory neurons (*mecDEG*). Adapted from Wang *et al.* 2017. **B)** Schematic of mechanosensory neurons with imaged regions highlighted (light gray boxes). **C)** Representative images of PLM axons for indicated genotypes visualized using P*mec-7*::RFP (*jsIs973*). Premature axon termination defects (arrows) occur when *mecDEG* targets GFP that is CRISPR engineered onto adhesome components (PAT-3::GFP, UNC-112::GFP and GFP::TLN-1). Vulva (asterisks) used as anatomical reference point for premature termination. **D)** Quantitation of premature axon termination defects in PLM neurons for indicated genotypes. Note premature axon termination occurs with *mecDEG* targeting of adhesome components and in *tln-1 (ok1648)* hypomorphic mutants. Means (bars) are shown for 5 or more counts (black dots) with 20 or more animals/count for each genotype. Error bars indicate SEM. Significance assessed using Student’s *t*-test with Bonferroni correction for multiple comparisons. ***p<0.001. Scale bar 10μm.

We chose to examine how *mecDEG*s targeting adhesome components affect PLM mechanosensory neurons because these neurons expressed PAT-3, UNC-112 and TLN-1 and display precise axon termination. Each PLM neuron has its soma in the tail of the animal and extends a single axon towards the midbody. Axon termination occurs with anatomic precision before the soma of the ALM mechanosensory neuron (**Fig 4B,C**). We began by evaluating several controls. First, we demonstrated that CRISPR tagging PAT-3, UNC-112 and TLN-1 with GFP (PAT-3::GFP, UNC-112::GFP and GFP::TLN-1) did not affect axon termination (**Fig 4D**). Second, we showed that expression of the *mecDEG* system alone did not affect axon termination (**Fig 4D**).

Next, we examined whether *mecDEG* co-expression with PAT-3::GFP, UNC-112::GFP or GFP::TLN-1 CRISPR affected axon termination. We observed premature axon termination defects, in which PLM axons terminated growth at or prior to the vulva, with *mecDEG* targeted degradation of PAT-3::GFP, UNC-112::GFP or GFP::TLN-1 CRISPR (**Fig 4C**). Quantitation indicated that premature termination defects were significant with cell-specific degradation of each adhesome component (**Fig 4D**). Moderate frequency of defects occurred with degradation of PAT-3::GFP CRISPR and defects were found at lower frequency when UNC-112::GFP CRISPR was impaired. Interestingly, degradation of GFP::TLN-1 resulted in a particularly high incidence of premature termination defects (**Fig 4D**). We note that TLN-1 has 7 isoforms, and our CRISPR strategy only GFP-tagged the largest isoform, isoform a, and a truncated isoform, isoform b (**Fig S4**). Thus, our results indicate that one or both of these isoforms mediate TLN-1 function during axon termination. The particularly high frequency of defects with GFP::TLN-1 CRISPR degradation could occur for several reasons. 1) TLN-1 is a primary adhesome signaling component in axon termination. 2) TLN-1 could have additional signaling activities outside the adhesome that influences axon termination. 3) Degradation of GFP::TLN-1 CRISPR by the *mecDEG* system could be more efficient than degradation of PAT-3::GFP or UNC-112::GFP.

To further evaluate the prominent premature axon termination phenotype observed when TLN-1 was impaired, we performed a transgenic rescue experiment. Here, we transgenically expressed TLN-1 using a pan-neuronal promoter in animals expressing both the *mecDEG* and GFP::TLN-1 CRISPR. We found that premature axon termination defects caused by degrading GFP::TLN-1 CRISPR were rescued by transgenic expression of untagged TLN-1, which cannot be degraded by *mecDEG* (**Fig 4C**). Quantitation of premature termination defects indicated that transgenic expression of TLN-1 lacking a GFP tag significantly reduced the frequency of defects (**Fig 4D**). To provide further genetic validation for TLN-1 affecting axon termination, we tested a *tln-1* deletion mutant, *(ok1648)*. We also observed significant premature termination defects in *tln-1 (ok1648)* mutants compared to wild-type animals (**Fig 4E**). However, the frequency of defects was significantly lower than *mecDEG* targeting of GFP::TLN-1 CRISPR.

Interestingly, our findings help us better understand the importance of TLN-1a and b isoforms in axon termination. *mecDEG* targeting GFP::TLN-1 CRISPR eliminates TLN-1a and b isoforms, which are the only isoforms that are N-terminally tagged with GFP (**Fig S4**). In contrast, the *tln-1 (ok1648)* deletion allele impairs several TLN-1 isoforms including the TLN-1a isoform, but not the TLN-1b isoform (**Fig S4**). Thus, our findings indicate that TLN-1a and b isoforms are particularly important for axon termination. Moreover, our observation that *tln-1 (ok1648)* mutants have lower frequency premature termination defects is likely to reflect that this allele is hypomorphic.

Collectively, our results support several conclusions. First, impairing any of the core adhesome components we tested, TLN-1/Talin, PAT-3/β-integrin and UNC-112/Kindlin, results in premature axon termination. Second, the cell-specific *mecDEG* strategy avoids lethality associated with null alleles and demonstrates that all three adhesome components function cell-autonomously in mechanosensory neurons to regulate axon termination. Finally, our results suggest that TLN-1 could be a particularly prominent player in axon termination.

### Adhesome components regulate axon termination by genetically opposing RPM-1

Our observation that perturbing core adhesome components in mechanosensory neurons resulted in premature axon termination prompted us to test genetic interactions between adhesome components and RPM-1. Previous studies demonstrated that RPM-1 functions cell-autonomously in mechanosensory neurons to regulate axon termination ^24, 25, 40^. In the PLM neurons, *rpm-1* protein null mutants ^24, 32^ result in failed axon termination defects where axons overgrow beyond the ALM cell body and hook towards the ventral side of the animal (**Fig 5A-C**).

To assess genetic interactions between multiple core adhesome components and RPM-1, we combined a *mecDEG* targeting PAT-3::GFP, UNC-112::GFP or GFP::TLN-1 CRISPR with an *rpm-1* null allele. As a control, we combined *rpm-1* mutants with the *mecDEG* system, which did not alter the frequency of failed axon termination defects caused by *rpm-1* loss of function (lf) (**Fig 5C**). Consistent with earlier experiments (**Fig 4D**), *mecDEG* targeting of PAT-3::GFP, UNC-112::GFP or GFP::TLN-1 CRISPR resulted in premature termination defects, the opposite defect to what occurs in *rpm-1* mutants (**Fig 5B, C**). Thus, opposing phenotypes suggest that RPM-1 and the adhesome could be functionally opposed during axon termination.

**Figure 5.**
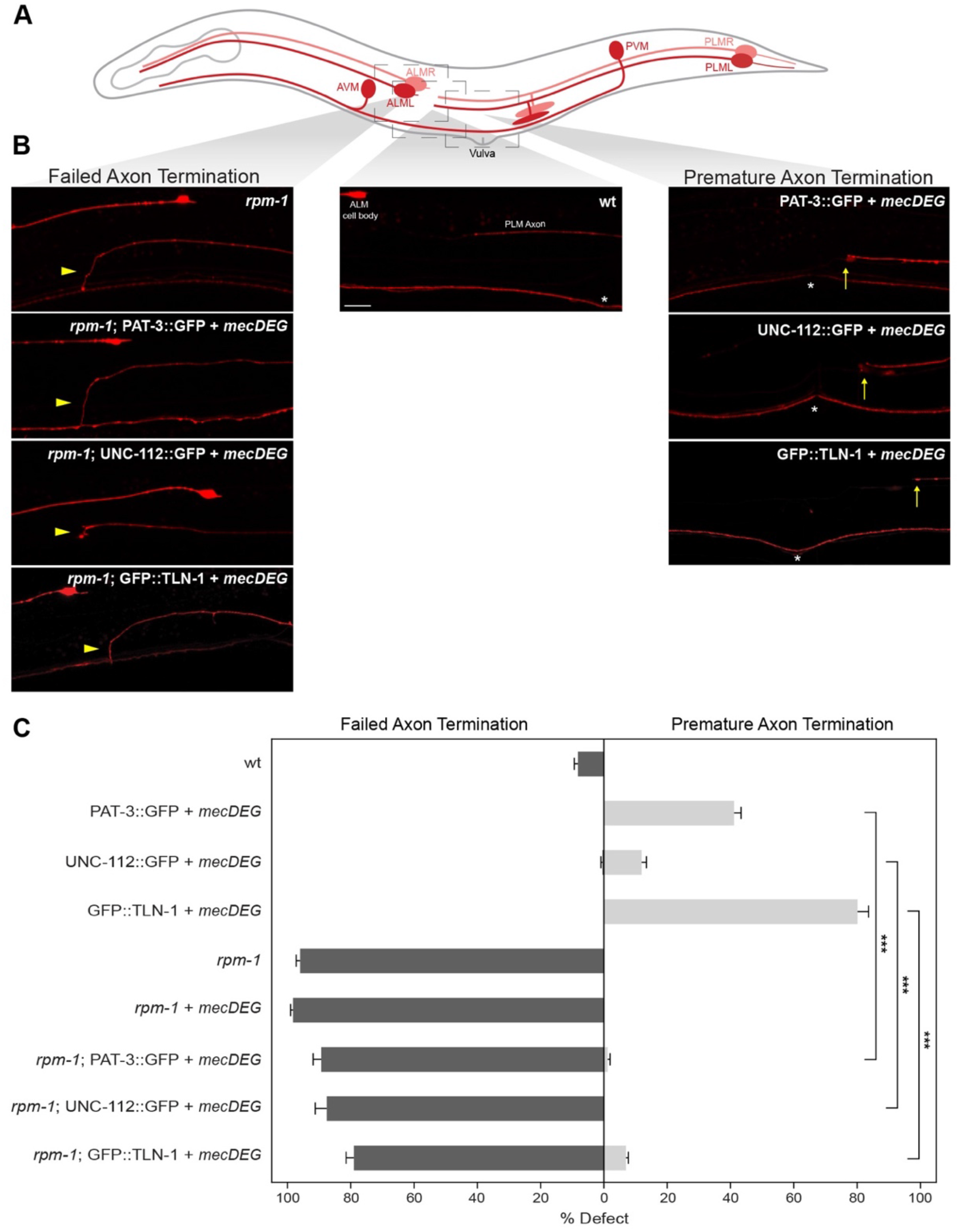
Opposing genetic interactions between adhesome components and *rpm-1* are required for accurate axon termination. **A)** Schematic of *C. elegans* mechanosensory neurons with imaged regions highlighted (light gray boxes). **B)** Representative images of PLM axons for indicated genotypes visualized using P*mec-7*::RFP (*jsIs973*). *rpm-1* mutants display failed axon termination defects (arrowhead) and *mecDEG* targeting of adhesome components (PAT-3::GFP, UNC-112::GFP and GFP::TLN-1 CRISPR) results in premature termination defects (arrows). *rpm-1* double mutants with adhesome component degradation (e.g. *rpm-1;* PAT-3::GFP + *mecDEG*) predominantly display failed termination defects. Vulva (asterisks) used as anatomical reference point for premature termination. **C)** Quantitation of failed axon termination defects (dark gray) and premature termination defects (light gray) in PLM neurons for indicated genotypes. Note premature termination defects are strongly suppressed in double mutants lacking *rpm-1* and adhesome components. Mean (bars) are shown for 5 or more counts (20 or more animals/count) for each genotype. Error bars indicate SEM. Significance assessed using Student’s *t-test* with Bonferroni correction. ***p<0.001. Scale bars 10μm.

In double mutants for *rpm-1* and all three adhesome components (e.g. *rpm-1;* PAT-3::GFP + *mecDEG*), we principally observed failed axon termination defects similar to *rpm-1* single mutants (**Fig 5B**). Quantitation indicated that premature termination defects primarily seen when adhesome components were impaired individually were substantially and significantly reduced in frequency in double mutants with *rpm-1* (**Fig 5C**). These results demonstrate that *rpm-1* genetically suppresses impaired adhesome function as a principal outcome. Thus, genetic interactions occur between multiple core adhesome components and *rpm-1*, which is consistent with our proteomic data indicating adhesome components physically associate with RPM-1.

### Talin and RPM-1 engage in a genetic ‘tug-of-war’ over the growth cone during axon termination

Our prior work demonstrated that RPM-1 regulates axon termination by promoting growth cone collapse in PLM mechanosensory neurons ^39^. Therefore, we performed developmental time-course experiments to examine whether genetic interactions between *rpm-1* and adhesome components affect growth cones during termination of axon outgrowth. We focused our developmental studies on Talin because *mecDEG* targeting of TLN-1 yielded the highest incidence of premature termination defects (**Fig 4**), suggesting it could be a key adhesome component involved in axon termination.

Prior developmental time-course studies solely with *rpm-1* mutants proved extremely challenging for several reasons ^39^. These experiments require precise synchronization of animal development, extensive imaging sessions, and evaluation of extremely small larval stage animals (**Fig 6A**, L1-L3). As a result, it was not technically feasible to perform developmental studies with all the control strains tested in axon termination studies on adults (**Fig 4D; 5C**). Therefore, we evaluated a more limited but essential set of controls. This included GFP::TLN-1 CRISPR alone as a negative control, because we observed no differences in axon termination for this control, *mecDEG* alone or wild-type animals (**Fig 4D**). We opted to evaluate *rpm-1* null mutants carrying the *mecDEG* control as we observed no differences in axon termination defects between this strain and *rpm-1* mutants lacking *mecDEG* (**Fig 5C**).

**Figure 6.**
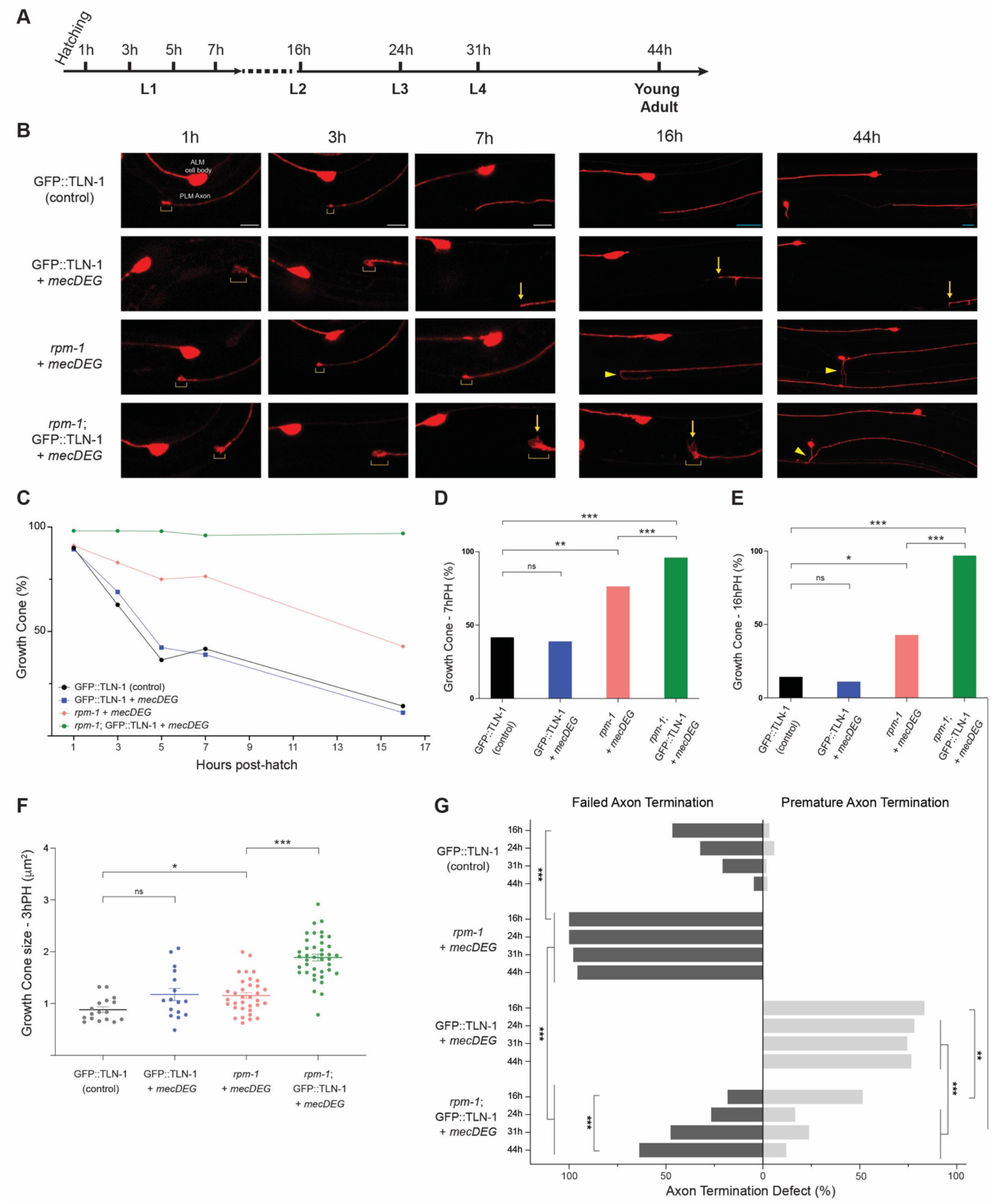
Developmental time-course studies demonstrate that *tln-1* and *rpm-1* affect axon termination by engaging in a genetic ‘tug-of-war’ over growth cones. **A)** Timeline of *C. elegans* larval development and key time points evaluated. **B)** Representative images of PLM axons for indicated genotypes visualized using P*mec-7*::RFP (*jsIs973*). Shown are axonal growth cones (brackets), premature termination sites (arrows) and failed termination defects (arrowheads). Note double mutants (*rpm-1;* GFP::TLN-1 + *mecDEG*) display failed growth cone collapse with persistent, enlarged growth cones. In double mutants, growth cones are initially in locations corresponding to premature axon termination (1 to 16h PH) but failed axon termination becomes the primary phenotype as development progresses (44h PH). Note AVM cell body is only visible at 44h PH on one side of the animal. **C)** Summary of quantitative results for growth cone frequency during development for indicated genotypes. **D-E)** Quantitative analysis of growth cone frequency at **D)** 7h PH and **E)** 16h PH. **F)** Quantitation of growth cone size at 3h PH for indicated genotypes. **G)** Quantitation of failed axon termination defects (dark gray) and premature termination defects (light gray) in PLM neurons for indicated genotypes at specified time points in development. For C-G, minimum of 21 PLM neurons scored for each time point and genotype. Mean (square, bar or line) are shown for each genotype. For F, dots represent a single animal. Error bars indicate SEM. For C-E and G, significance assessed using Fisher’s exact test. For F, significance assessed using Student’s *t-test* with Bonferroni correction for growth cone size. ns=not significant, *p<0.05, **p<0.01, ***p<0.001. Scale bars are 5μm (1-7h PH, white bars) or 10μm (16-44h PH, teal bars).

We began by evaluating growth cone frequency and size at different developmental time points across all larval stages - L1, L2, L3 and L4 (**Fig 6A**). Several observations were consistent with prior studies. Growth cone frequency decreased between 1h and 16h post-hatch (PH) in control GFP::TLN-1 CRISPR animals (**Fig 6B-E**). This occurred with similar timelines and decreasing growth cone frequency as previously observed for wild-type animals ^39^. Decreasing growth cone frequency was accompanied by termination of axon outgrowth which occurred between 7 and 16h PH (**Fig 6B, G**). Based on our scoring criteria, failed termination defects occurred in GFP::TLN-1 controls at moderate frequency at 16h PH. However, this reflects a normal part of development, as overextension is alleviated over time when animals increase in size and form gaps between ALM cell bodies and PLM axon termination sites ^39, 44^. Thus, overextension present at this time point is not due to CRISPR engineering GFP onto TLN-1.

In *rpm-1* single mutants carrying *mecDEG* alone control (*rpm-1* + *mecDEG*), growth cone collapse was impaired resulting in both increased growth cone frequency (**Fig 6B-E**) and increased growth cone size (**Fig 6B, F**) compared to GFP::TLN-1 controls. Due to impaired growth cone collapse, failed axon termination defects emerge at high frequency in *rpm-1* + *mecDEG* mutants by 16h PH (L2) and persist through young adult animals (44h PH) (**Fig 6B, G**). Our findings with *rpm-1* + *mecDEG* are consistent with prior findings with *rpm-1* mutants ^39^. Our developmental findings are also consistent with the incidence of failed termination defects being similar in *rpm-1* mutants and *rpm-1* + *mecDEG* mutants (**Fig 5C**). Taken together, these results indicate that the *mecDEG* system does not further influence growth cone development beyond the effects of *rpm-1* (lf) in the absence of an endogenous CRISPR-tagged GFP target protein.

We then evaluated GFP::TLN-1 + *mecDEG* animals (*tln-1* single mutants), and tested the hypothesis that early/premature growth cone collapse results in premature axon termination defects. Our results disproved this hypothesis, as GFP::TLN-1 + *mecDEG* did not have altered growth cone frequency (**Fig6 B-E**) or growth cone size (**Fig 6B, F**) when compared to GFP::TLN-1 CRISPR control animals. Interestingly, GFP::TLN-1 + *mecDEG* animals displayed growth cones in locations corresponding with premature termination as early as 1h PH (**Fig 6B**), and premature termination defects accumulated between 7 and 16h PH (**Fig 6B, G**). These findings suggest that premature axon termination defects in TLN-1 mutants occur due to reduced axon extension.

Next, we evaluated *rpm-1*; GFP::TLN-1 + *mecDEG* animals, which are double mutants for *rpm-1* and *tln-1*. We observed enhanced growth cone frequency (**Fig 6B-E**) and increased growth cone size (**Fig 6B, F**) in *rpm-1*; GFP::TLN-1 + *mecDEG* double mutants compared to *rpm-1* + *mecDEG* single mutants. Growth cones also persisted at very high frequency out to 16h PH (**Fig 6B, C, E**). This is dramatically beyond the developmental time point where growth cones collapse (**Fig 6B-E**) and axon termination occurs in control GFP::TLN-1 animals (**Fig 6B, G**) or wild-type animals ^39^. It is possible that in the double mutant, without RPM-1 driving the growth cone to collapse and without TLN-1 promoting axon extension, altering opposing genetic forces results in a higher incidence of persistent, larger growth cones. Interestingly, growth cones that persist at 16h PH in *rpm-1*; GFP::TLN-1 + *mecDEG* double mutants present as a mixture of potential failed and premature termination events based on anatomical locations where growth cones are present at this time point (**Fig 6B, G**). While axon termination has not occurred at 16hr PH in double mutants, we applied the same scoring scheme used to score axon termination in adults. As development progresses, potential premature termination in *rpm-1*; GFP::TLN-1 + *mecDEG* double mutants is overcome and the failed termination phenotype present in *rpm-1* single mutants primarily takes over between 24 and 44h PH (**Fig 6B, G**).

Interestingly, our findings suggest that failed growth cone collapse from *rpm-1* (lf) and reduced axon extension from *tln-1* (lf) initially results in shorter axons with enlarged growth cones early in development of double mutants. Over time, there is a functional and genetic ‘tug-of-war’ on growth cone development between the TLN-1 adhesome component and RPM-1. Ultimately, defects caused by *rpm-1* (lf) become dominant and defects caused by GFP::TLN-1 + *mecDEG* are suppressed. These findings indicate that TLN-1 and RPM-1 control axon termination via an opposing genetic interface that is important for proper growth cone development and accurate, timely termination of axon outgrowth.

## DISCUSSION

Recent bioinformatic and proteomic studies have identified an enormous integrin signaling network called the adhesome, which is evolutionarily conserved from humans through *C. elegans* ^8, 38^. Thus, *C. elegans* is an ideal model to evaluate the composition and function of a simplified adhesome, and allows us to test the functional relationship between the adhesome and prominent players in nervous system development. Three core components of the putative *C. elegans* adhesome are PAT-3/β-integrin, UNC-112/Kindlin and TLN-1/Talin. Prior studies have principally examined their roles in body wall muscle development, which leads to lethality in null mutants ^42, 45^. Much less is known about how these adhesome components influence nervous system development in *C. elegans*. A small number of studies have used hypomorphic alleles to show that integrins regulate axon guidance in motor neurons ^12, 46, 47^. At present, we know particularly little about how UNC-112/Kindlin and TLN-1/Talin shape axon development in vivo.

Here, we have taken a first step towards obtaining a more comprehensive, mechanistic understanding of how the adhesome shapes axonal and nervous system development. To do so, we integrated proteomics with cell-specific genetic tools to evaluate how multiple core adhesome components shape axon development in the mechanosensory neurons of *C. elegans*. CRISPR engineering and targetted degradation of TLN-1, UNC-112 and the PAT-3 circumvented lethality caused by global null alleles ^42^. Our results demonstrate that these core adhesome components are expressed (**Fig 3**) and function cell-autonomously within mechanosensory neurons to regulate termination of axon outgrowth (**Fig 4**). Moreover, our proteomic and genetic results indicate that a subset of adhesome components regulate axon development by physically associating with (**Fig 1, 2; Table 1**) and functionally opposing the RPM-1 ubiquitin ligase signaling hub (**Fig 5**). Our findings now provide orthogonal, unbiased identification and validation of the adhesome using proteomic, genetic and developmental studies in a whole organism setting.

Developmental studies were instrumental in evaluating the cellular mechanism underpinning the genetic relationship between the adhesome and RPM-1. Our results indicate that genetic opposition between TLN-1 and RPM-1 occurs because these players converge on developing axons in a functional ‘tug-of-war’ (**Fig 6**). Interestingly, the opposing genetic interface we have identified is required to integrate axon extension with growth cone collapse to yield spatially and temporally accurate axon termination. Our results indicate that the core adhesome components TLN-1, UNC-112 and PAT-3 promote axon growth, as axons do not grow as far and terminate prematurely when these components are impaired compared to CRISPR engineered GFP controls (**Fig 3, 6**) or wild-type animals (**Fig 3**). Our findings with PAT-3 in mechanosensory neurons are consistent with prior studies indicating β-integrins can regulate axon outgrowth in cultured neurons and axons of the corpus callosum in rodents ^16, 48^. Our observation that UNC-112 is required for axon extension and termination of outgrowth in a whole animal setting expands upon prior studies that used shRNA to knockdown Kindlin in cultured neurons ^20, 49^. The third core adhesome component we tested, TLN-1, has not been previously shown to shape axon development in any system. Our observation that premature termination defects occur with higher frequency when TLN-1 is perturbed compared to UNC-112 suggests that Talin could be a more prominent adhesome component in the axon termination process. This supports recent commentary, based on studies outside the nervous system, that Talin could be a master player in adhesome signaling ^17^. However, it is important to also note that our results indicate Talin and Kindlin function are both needed for accurate, efficient axon termination.

In contrast to premature axon termination that occurs when adhesome components are perturbed, loss of RPM-1 presents the opposing phenotype of failed axon termination and overgrowth due to impaired growth cone collapse (**Fig 3, 6**) ^39^. In double mutants between RPM-1 and core adhesome components including TLN-1, PAT-3 or UNC-112, we observed genetic suppression of premature termination defects by adulthood. While this potentially suggests adhesome components could inhibit RPM-1, we note this tentatively because we could not use null alleles for these adhesome components due to issues with viability. However, our results do conclusively demonstrate that when both a core adhesome component and RPM-1 are perturbed there is an imbalance in genetic forces affecting axon termination. Interestingly, developmental time-course studies revealed that when both TLN-1 and RPM-1 are impaired we initially observe compounded increases in growth cone frequency and growth cone size (**Fig 6B-F**). While this might appear counter-intuitive, we think the opposing genetic effects caused by impairing TLN-1 and RPM-1 yield increased growth cone size because no outcome on axon growth is realized early in development. Overtime, impaired RPM-1 signaling and failed growth cone collapse takes over with failed axon termination and overgrowth increasing through development, and eventually becoming the primary phenotypic outcome. Consistent with this model, we observe growth cone persistence in double mutants of TLN-1 and RPM-1 for many hours beyond when growth cone collapse occurs and axon termination is complete in control GFP::TLN-1 CRISPR animals (**Fig 6B-G**) or wild-type animals (**Fig 3**) ^39^. To our knowledge, this is one of the most striking examples of impaired growth cone collapse that has been observed in vivo. Thus, we have identified a prominent genetic interface between multiple adhesome components and RPM-1 signaling that is required to integrate axon extension and growth cone collapse during termination of axon outgrowth.

Interestingly, our findings have revealed further links between RPM-1 signaling and components of the adhesome. One example is the Cdk5 kinase which phosphorylates Talin and is inhibited by RPM-1 ubiquitin ligase activity ^34, 50^. The MIG-15/NIK kinase interacts genetically with RPM-1 and physically binds PAT-3 β-Integrin ^46, 51^. Integrins and Kindlin affect axon regeneration and axon degeneration, functional contexts that also involve RPM-1 and its murine ortholog Phr1 ^4, 27, 52–55^. Finally, studies in mice indicate that loss of Phr1 or β-Integrin result in developmental defects in the corpus callosum ^48, 56, 57^. Our results now provide genetic and developmental mechanisms that potentially explain these previously unrecognized links between core adhesome components and RPM-1 signaling.

There is growing evidence that integrin signaling is involved in brain disorders ^2, 3^. By combining proteomics and computational network analysis, we identified human genetic variants associated with neurobehavioral abnormalities (*i.e.* intellectual disability, autism or developmental delay) in 75% of adhesome components that interact with RPM-1 (**Fig 2**). This is a particularly interesting observation, as a recent study identified multiple de novo variants in *RPM-1/MYCBP2* that were shown to be deleterious by CRISPR editing in *C. elegans* ^32^. Patients with *MYCBP2* variants have a neurodevelopmental disorder called MDCD that features corpus callosum defects and a spectrum of neurobehavioral deficits including developmental delay, intellectual disability and autism. Thus, we have identified a subnetwork of adhesome components that are associated with neurobehavioral abnormalities, and that interact physically and functionally with a gene that causes a neurodevelopmental disorder. Our findings now point to the concept of adhesome biology as a potentially rich direction for future studies on nervous system development and disease.

## METHODS

### Genetics and strains

*C. elegans* N2 isolate was used for all experiments. Animals were maintained using standard procedures. The following mutant alleles were used: *rpm-1*(*ju44*) V and *tln-1*(*ok1648*) I. The following integrated transgene was used: *jsIs973* [P_mec-7_::mRFP] III. The following MosSCI transgene was used: *itSi953* [P_mec-18_::vhhGFP4::ZIF-1::operon-linker::mKate::tbb-2 3’UTR + Cbr-unc-119(+)] II, which we refer to as *mecDEG*. All MosSCI transgenes were inserted into *ttTi5605* on LG II. The following CRISPR alleles were used: *unc-112*(*bgg68* [UNC-112::GFP CRISPR]), *pat-3*(*bgg86* [PAT-3::GFP CRISPR]), *tln-1*(*zh117* [GFP::TLN-1 CRISPR]). All transgenes and CRISPR alleles used for specific experiments are described in Table S4. All mutants and transgenic lines were outcrossed four or more times prior to experiments. Animals were grown at 23°C for genetic analysis. Sequences of all genotyping primers for mutants and CRISPR strains can be found in Table S5.

### Molecular biology and Transgenics

Four fragments of the *tln-1* genomic DNA (gDNA) (pBG-1088) were amplified from N2 genomic DNA using iProof High-Fidelity DNA polymerase (BioRad) or Q5® High-Fidelity DNA Polymerase (NEB). Fragments were then assembled into a single *tln-1* gDNA clone in a pCR8 vector using HiFi DNA assembly (NEB). Using gateway technology (LR clonase II, Invitrogen), *tln-1* gDNA was recombined with a plasmid containing P_rgef-1_::FLAG and *unc-54* 3’UTR (PBG-GY134) to generate the final P_rgef-1_FLAG::*tln-1* plasmid (pBG-GY1100) used for microinjection to generate the TLN-1 rescue strain *bggEx172* (P_rgef-1_FLAG::TLN-1). All constructs were fully sequenced.

Transgenic extrachromosomal arrays were generated using standard microinjection procedures for *C. elegans*. Injection conditions and transgene construction details are specified in Table S6.

### CRISPR/Cas9 engineering

CRISPR alleles were engineered utilizing *dpy-10* co-CRISPR and direct injection of *Cas9* ribonucleoprotein complexes. Injection mixes contained tracrRNA (IDT), repair template (PCR or ssODN (Ultramers DNA oligo IDT)), and recombinant Cas9 protein purified from Rosetta 2 *E. coli* (Millipore EMD, #71397). Ribonucleoprotein complexes were placed at 37°C for 15 minutes prior to injection. Gene edits were confirmed by sequencing, and edited strains were outcrossed four or more times to N2 animals. All CRISPR injection conditions are shown in Table S6. All crRNA and repair template sequences are shown in Table S7.

### C. elegans proteomics

Detailed methodology for *C. elegans* AP-proteomics was previously described in detail ^25, 33^. In brief, experiments were conducted on mixed-stage animals transgenically expressing GS::RPM-1, GS::RPM-1 LD substrate ‘trap’ or GS::GFP (negative control). Animals were harvested, separated from culture debris by sucrose flotation centrifugation, frozen in liquid nitrogen, and ground into submicron particles using a cryomill (Retsch). Samples were then lysed under various detergent conditions. Resulting whole-animal lysates were centrifuged to separate particulate material. Lysates were incubated with IgG-coupled Dynabeads (80 mg total protein with 500uL of beads) at 4°C for 4 hours. Samples were run on Tris-Glycine SDS-PAGE gels, followed by Coomassie staining and in-gel trypsin digested. The resulting peptide pools were acidified, desalted, dried, and subsequently resuspended in 100 μl of 0.1% formic acid. Samples were run on an Orbitrap Fusion Tribrid Mass Spectrometer (ThermoFisher Scientific) coupled to an EASY-nLC 1000 system and on-line eluted on an analytical RP column (0.075 × 250 mm Acclaim PepMap RLSC nano Viper, ThermoFisher Scientific). Scaffold (Proteome Software) was used to analyze MS/MS data to identify peptides and proteins in samples. All control and test samples for single proteomics experiments were blinded for genotype and underwent mass spectrometry together.

We performed a total of 7 independent AP-proteomics experiments with RPM-1. Identification of adhesome components physically associated with RPM-1 was based on three criteria: 1) Detection in one or more proteomic experiments. 2) 1.5x or greater total spectra enrichment in GS::RPM-1 or GS::RPM-1 LD sample compared to GS::GFP negative control. 3) Elimination of ribosomal and vitellogenin proteins, as they are widely recognized as proteomic contaminants.

### Analysis of axon termination

We evaluated axon termination in PLM neurons using the transgene *jsIs973* [P_mec-7_::mRFP]. We considered three termination zones based on anatomical criteria when scoring animals. 1) Wild-type axon termination was considered to occur after the vulva but before the ALM soma. 2) Failed axon termination defects occurred when axons overgrew and overextended past the ALM soma and/or formed ventral ‘hooks’ that occurred after the vulva or after the ALM cell body. 3) Premature termination defects occurred if the axon terminated growth at or before the vulva.

To quantify axon termination, young adult animals were anesthetized using 10 μM levamisole in M9 buffer. Animals were mounted onto 2% agar pads on glass slides with coverslips. Animals were visualized using a Leica DM5000 B (CTR5000) epifluorescent microscope equipped with a 40x oil-immersion objective.

For image acquisition, young adult animals were anesthetized using either 10 μM levamisole or a 3% (v/v) solution of 1-phenoxy-2-propanol in M9 buffer. Animals were mounted on 3% agarose pads on glass slides with coverslips. Images were captured using a Zeiss LSM 710 equipped with 40x oil-immersion objective.

### Developmental analysis

The transgene *jsIs973* (P_mec-7_::mRFP) was used to visualize growth cone frequency and growth cone size in PLM neurons and to measure the distance between PLM axon termination sites and ALM somas.

Developmental time-course studies were performed as previously described ^39^. Animals were synchronized by collecting freshly hatched L1 larvae every 10-15 minutes and culturing on plates to desired time points at 23°C. At select time points between 1 and 44 h PH, animals were mounted in 5μM levamisole (1-16h PH) or 10μM levamisole (24-44h PH) in M9 buffer on 3% agarose pads. Images were acquired under 63× (1-31h PH) or 40× (44h PH) magnification on a Zeiss LSM 710 laser scanning confocal microscope.

Image analysis of growth cone width and area as well as growth cone distance from the ALM soma was executed using Fiji/ImageJ software from NIH image. We defined growth cones compared to terminated axon tips using both morphological assessment and quantitative measurements of the ratio of growth cone width to axon width as previously described ^39^. A growth cone was defined as having a ratio of growth cone width to axon width of 1.5x or greater. Images of all genotypes and time points were blinded and mixed before analysis using custom software run with Python. We used criteria above to categorize termination defects in adults (44h PH) when a vulva could be observed and used as an anatomical reference point. For L1-L4 larval stages (16h to 31h PH) where a vulval was not present, we defined axon termination defects (failed or premature) by measuring the distance between the middle of the ALM soma to the end of the PLM axon (growth cone or terminated axon tip). Premature termination was classified depending on the distance between ALM and terminated axon tip or growth cone during larval stages as follows: 15μm for 1, 3, 5, 7h PH, 24μm for 16h PH, 55μm for 24h PH, 65μm for 31h PH. These metrics were derived based on control animals for a given time point. Failed termination was defined as an axon terminating anterior to the ALM soma or the presentation of a ‘hook’ towards the ventral cord for all stages.

### AlphaFold prediction

Structures of GFP fusion proteins were predicted using the LocalColabFold version of ColabFold 1.5.0 using pdb templates with “--templates” and relaxed using “--amber” ^58^. Five models were predicted for each protein and the model with the highest predicted local distance difference test (plDDT) was presented for each protein. Predicted structures were adjusted and colorized in PyMOL (PyMOL Molecular Graphics System, Version 2.5.0, Schrödinger, LLC). The initial output for GFP::TLN-1 resembled one of its templates: the closed, autoinhibited conformation of human TALIN 1 (PDB 6R9T). For visual clarity, overlapping domains were manually separated by adjusting dihedral angles in residues with pLDDT scores equal to 50 or less and lacking secondary structure. Thus, the presented model represents a partially open conformation where the head and tail regions show their predicted compact forms but do not interact. PAT-3::GFP was also similarly adjusted to separate the intracellular and extracellular domains.

### *C. elegans* integrin adhesome computational network analysis

Based on a list of 232 molecular components that comprise the mammalian adhesome network ^8, 9^, we sought to identify all potential *C. elegans* orthologs based on three primary criteria: 1) 30% or higher sequence similarity 2) 20% or higher sequence identity, and 3) prior annotation in the

*C. elegans* adhesome ^38^, which we carefully re-evaluated here ourselves. We determined that 132 mammalian adhesome components have *C. elegans* orthologs, and of these 123 mammalian components have unique, single orthologous components in *C. elegans*.

Utilizing STRING software ^59^, these 123 conserved, unique *C. elegans* integrin adhesome components were mapped into a computational network of predicted protein-protein interactions. Line connections between proteins indicate prior evidence for protein-protein interactions. Predicted protein-protein interactions were based on experiments and/or databases, and we reported outcomes with a minimum interaction score of 0.4.

RPM-1 AP-proteomics data was overlaid onto the predicted *C. elegans* integrin adhesome signaling network. Any orthologous integrin adhesome protein detected in RPM-1 AP-proteomics are shown in blue. Their overall enrichment in GS::RPM-1 test samples over GS::GFP negative control samples are represented by the size of the circle with increasing size indicating greater significance.

### Statistical analysis

#### Proteomics

For AP-proteomics, protein hits that were identified with 1.5x or greater enrichment in GS::RPM-1 or GS::RPM-1 LD test samples over GS::GFP negative control samples were subjected to unpaired, two-tailed Student’s *t*-test to assess significance using total spectral counts across seven independent proteomic experiments. Each experiment was considered an individual n for analysis. For statistical comparisons in Table 1, we normalized total spectral counts for the candidate hit protein to both its molecular weight as well as the total spectral counts and molecular weight of the affinity purification target, as previously detailed ^33^. Similar normalization for protein size was done for calculation of fold enrichment of candidates between GS::RPM-1 and GS::RPM-1 LD samples in Table 1.

#### Axon termination

For analysis of PLM axon termination, statistical comparisons were performed using unpaired, two-tailed Student’s *t*-test with Bonferroni correction for multiple comparisons. Statistical analysis was performed using GraphPad Prism software. Error bars are SEM. Significance was defined as p < 0.05. Histograms represent averages from 4 to 10 counts (25–35 neurons/count) for each genotype from three or more independent experiments. Dots in plots represent averages from single-counting sessions.

#### Developmental analysis

For growth cone frequency in PLM neurons, statistical comparisons were made by Fisher’s exact test using GraphPad Prism software. Significance was defined as p < 0.05. Histograms represent averages from at least 21 animals obtained from 2 or more, independent imaging session for each genotype.

For growth cone size in PLM neurons, statistical comparisons were made by Student’s *t*-test with Bonferroni correction for multiple comparisons using GraphPad Prism software. Error bars are SEM. Significance was defined as p < 0.05. Lines in scatter plots represent averages from at least 16 animals obtained from 2 or more, independent imaging session for each genotype. Dots shown in plots represent values from each individual animal.

For axon termination in PLM neurons, statistical comparisons were made by Fisher’s exact test using GraphPad Prism software. Significance was defined as p < 0.05. Histograms represent averages from at least 23 animals obtained from 2 or more, independent imaging session for each genotype.

## ACKNOWLEDGEMENTS

We would like to thank Dr. George Tsaprailis from the Proteomics Core at the University of Florida Scripps Labs (formerly The Scripps Research Institute – Florida) for his aid in analysis of mass spectra. We thank the *C. elegans* Genetics Center for strains and the Wormbase genetic resource database. B.G. was supported by National Institutes of Health Grant R01 NS072129. J.A. was supported by NINDS Diversity Supplement: R01 NS072129-S1. J.S.P. was supported by the NIDA Diversity Supplement: R01 DA048036-S1.

## AUTHOR CONTRIBUTION

J.A. performed *C. elegans* genetics, data acquisition, and data analysis for axon termination and adhesome component expression. He also performed analysis of RPM-1 AP proteomics data, computational analysis of the *C. elegans* adhesome network and identified genetic links between adhesome components and human neurobehavioral abnormalities. M.D. performed and analysed developmental time-course experiments including confocal microscopy, provided input on experimental design, and co-supervised J.A. K.J.O. performed proteomics, identified an initial cohort of adhesome components in RPM-1 proteomics data, and provided intellectual input and initial testing of the *mecDEG* system. J.S.P performed AlphaFold predictions of GFP tagged adhesome components and generated predicted structural models. N.T.N. made the TLN-1 rescue plasmid. K.D-G performed confocal microscopy and data acquisition for adhesome component expression. E.C. executed experiments with the *tln-1* hypomorphic allele. M.A.B provided expertise and input on developmental time-course experiments and provided conceptual input on the manuscript. B.G. provided input on experimental design, provided supervision for all personnel, and managed project funding. B.G., M.D. and J.A. wrote the manuscript.

## REFERENCES

1. Myers, J.P., Santiago-Medina, M., and Gomez, T.M. (2011). Regulation of axonal outgrowth and pathfinding by integrin-ECM interactions. Dev Neurobiol 71, 901–923. 10.1002/dneu.20931.

2. Park, Y.K., and Goda, Y. (2016). Integrins in synapse regulation. Nat Rev Neurosci 17, 745–756. 10.1038/nrn.2016.138.

3. Lilja, J., and Ivaska, J. (2018). Integrin activity in neuronal connectivity. J Cell Sci 131. 10.1242/jcs.212803.

4. Eva, R., and Fawcett, J. (2014). Integrin signalling and traffic during axon growth and regeneration. Curr Opin Neurobiol 27, 179–185. 10.1016/j.conb.2014.03.018.

5. Wickstrom, S.A., Radovanac, K., and Fassler, R. (2011). Genetic analyses of integrin signaling. Cold Spring Harb Perspect Biol 3. 10.1101/cshperspect.a005116.

6. Abram, C.L., and Lowell, C.A. (2009). The ins and outs of leukocyte integrin signaling. Annu Rev Immunol 27, 339–362. 10.1146/annurev.immunol.021908.132554.

7. Cooper, J., and Giancotti, F.G. (2019). Integrin Signaling in Cancer: Mechanotransduction, Stemness, Epithelial Plasticity, and Therapeutic Resistance. Cancer cell 35, 347–367. 10.1016/j.ccell.2019.01.007.

8. Winograd-Katz, S.E., Fassler, R., Geiger, B., and Legate, K.R. (2014). The integrin adhesome: from genes and proteins to human disease. Nat Rev Mol Cell Biol 15, 273–288. 10.1038/nrm3769.

9. Zaidel-Bar, R., Itzkovitz, S., Ma’ayan, A., Iyengar, R., and Geiger, B. (2007). Functional atlas of the integrin adhesome. Nat Cell Biol 9, 858–867. 10.1038/ncb0807-858.

10. Schiller, H.B., Friedel, C.C., Boulegue, C., and Fassler, R. (2011). Quantitative proteomics of the integrin adhesome show a myosin II-dependent recruitment of LIM domain proteins. EMBO reports 12, 259–266. 10.1038/embor.2011.5.

11. Horton, E.R., Byron, A., Askari, J.A., Ng, D.H.J., Millon-Fremillon, A., Robertson, J., Koper, E.J., Paul, N.R., Warwood, S., Knight, D., et al. (2015). Definition of a consensus integrin adhesome and its dynamics during adhesion complex assembly and disassembly. Nat Cell Biol 17, 1577–1587. 10.1038/ncb3257.

12. Baum, P.D., and Garriga, G. (1997). Neuronal migrations and axon fasciculation are disrupted in ina-1 integrin mutants. Neuron 19, 51–62.

13. Condic, M.L., and Letourneau, P.C. (1997). Ligand-induced changes in integrin expression regulate neuronal adhesion and neurite outgrowth. Nature 389, 852–856. 10.1038/39878.

14. Pasterkamp, R.J., Peschon, J.J., Spriggs, M.K., and Kolodkin, A.L. (2003). Semaphorin 7A promotes axon outgrowth through integrins and MAPKs. Nature 424, 398–405. 10.1038/nature01790.

15. Robles, E., and Gomez, T.M. (2006). Focal adhesion kinase signaling at sites of integrin-mediated adhesion controls axon pathfinding. Nat Neurosci 9, 1274–1283. 10.1038/nn1762.

16. Gupton, S.L., and Gertler, F.B. (2010). Integrin signaling switches the cytoskeletal and exocytic machinery that drives neuritogenesis. Dev Cell 18, 725–736. 10.1016/j.devcel.2010.02.017.

17. Klapholz, B., and Brown, N.H. (2017). Talin - the master of integrin adhesions. J Cell Sci 130, 2435–2446. 10.1242/jcs.190991.

18. Rognoni, E., Ruppert, R., and Fassler, R. (2016). The kindlin family: functions, signaling properties and implications for human disease. J Cell Sci 129, 17–27. 10.1242/jcs.161190.

19. Kerstein, P.C., Jacques-Fricke, B.T., Rengifo, J., Mogen, B.J., Williams, J.C., Gottlieb, P.A., Sachs, F., and Gomez, T.M. (2013). Mechanosensitive TRPC1 channels promote calpain proteolysis of talin to regulate spinal axon outgrowth. J Neurosci 33, 273–285. 10.1523/JNEUROSCI.2142-12.2013.

20. Tan, C.L., Andrews, M.R., Kwok, J.C., Heintz, T.G., Gumy, L.F., Fassler, R., and Fawcett, J.W. (2012). Kindlin-1 enhances axon growth on inhibitory chondroitin sulfate proteoglycans and promotes sensory axon regeneration. J Neurosci 32, 7325–7335. 10.1523/JNEUROSCI.5472-11.2012.

21. Grueber, W.B., and Sagasti, A. (2010). Self-avoidance and tiling: Mechanisms of dendrite and axon spacing. Cold Spring Harb Perspect Biol 2, a001750. 10.1101/cshperspect.a001750.

22. Harris, J.A., Mihalas, S., Hirokawa, K.E., Whitesell, J.D., Choi, H., Bernard, A., Bohn, P., Caldejon, S., Casal, L., Cho, A., et al. (2019). Hierarchical organization of cortical and thalamic connectivity. Nature 575, 195–202. 10.1038/s41586-019-1716-z.

23. Grill, B., Murphey, R.K., and Borgen, M.A. (2016). The PHR proteins: intracellular signaling hubs in neuronal development and axon degeneration. Neural Dev 11, 8. 10.1186/s13064-016-0063-0.

24. Schaefer, A.M., Hadwiger, G.D., and Nonet, M.L. (2000). rpm-1, a conserved neuronal gene that regulates targeting and synaptogenesis in C. elegans. Neuron 26, 345–356.

25. Crawley, O., Opperman, K.J., Desbois, M., Adrados, I., Borgen, M.A., Giles, A.C., Duckett, D.R., and Grill, B. (2019). Autophagy is inhibited by ubiquitin ligase activity in the nervous system. Nature communications 10, 5017. 10.1038/s41467-019-12804-3.

26. Park, E.C., and Rongo, C. (2018). RPM-1 and DLK-1 regulate pioneer axon outgrowth by controlling Wnt signaling. Development 145. 10.1242/dev.164897.

27. Virdee, S. (2022). An atypical ubiquitin ligase at the heart of neural development and programmed axon degeneration. Neural Regen Res 17, 2347–2350. 10.4103/1673-5374.338992.

28. Pao, K.C., Wood, N.T., Knebel, A., Rafie, K., Stanley, M., Mabbitt, P.D., Sundaramoorthy, R., Hofmann, K., van Aalten, D.M.F., and Virdee, S. (2018). Activity-based E3 ligase profiling uncovers an E3 ligase with esterification activity. Nature 556, 381–385. 10.1038/s41586-018-0026-1.

29. Desbois, M., Crawley, O., Evans, P.R., Baker, S.T., Masuho, I., Yasuda, R., and Grill, B. (2018). PAM forms an atypical SCF ubiquitin ligase complex that ubiquitinates and degrades NMNAT2. J Biol Chem 293, 13897–13909. 10.1074/jbc.RA118.002176.

30. Lewcock, J.W., Genoud, N., Lettieri, K., and Pfaff, S.L. (2007). The ubiquitin ligase Phr1 regulates axon outgrowth through modulation of microtubule dynamics. Neuron 56, 604–620.

31. Chang, C., Banerjee, S.L., Park, S.S., Zhang, X., Cotnoir-White, D., Desbois, M., Grill, B., and Kania, A. (2023). Ubiquitin ligase and signalling hub MYCBP2 is required for efficient EPHB2 tyrosine kinase receptor function. bioRxiv. 10.1101/2023.06.12.544638.

32. AlAbdi, L., Desbois, M., Rusnac, D.V., Sulaiman, R.A., Rosenfeld, J.A., Lalani, S., Murdock, D.R., Burrage, L.C., Undiagnosed Diseases, N., Billie Au, P.Y., et al. (2023). Loss-of-function variants in MYCBP2 cause neurobehavioural phenotypes and corpus callosum defects. Brain 146, 1373–1387. 10.1093/brain/awac364.

33. Desbois, M., Pak, J.S., Opperman, K.J., Giles, A.C., and Grill, B. (2023). Optimized protocol for in vivo affinity purification proteomics and biochemistry using C. elegans. STAR Protoc 4, 102262. 10.1016/j.xpro.2023.102262.

34. Desbois, M., Opperman, K.J., Amezquita, J., Gaglio, G., Crawley, O., and Grill, B. (2022). Ubiquitin ligase activity inhibits Cdk5 to control axon termination. PLoS genetics 18, e1010152. 10.1371/journal.pgen.1010152.

35. Baker, S.T., Opperman, K.J., Tulgren, E.D., Turgeon, S.M., Bienvenut, W., and Grill, B. (2014). RPM-1 Uses Both Ubiquitin Ligase and Phosphatase-Based Mechanisms to Regulate DLK-1 during Neuronal Development. PLoS genetics 10, e1004297. 10.1371/journal.pgen.1004297.

36. Grill, B., Bienvenut, W.V., Brown, H.M., Ackley, B.D., Quadroni, M., and Jin, Y. (2007). C. elegans RPM-1 Regulates Axon Termination and Synaptogenesis through the Rab GEF GLO-4 and the Rab GTPase GLO-1. Neuron 55, 587–601.

37. Grill, B., Chen, L., Tulgren, E.D., Baker, S.T., Bienvenut, W., Anderson, M., Quadroni, M., Jin, Y., and Garner, C.C. (2012). RAE-1, a Novel PHR Binding Protein, Is Required for Axon Termination and Synapse Formation in Caenorhabditis elegans. J Neurosci 32, 2628–2636. 32/8/2628 [pii]10.1523/JNEUROSCI.2901-11.2012.

38. Zaidel-Bar, R. (2009). Evolution of complexity in the integrin adhesome. J Cell Biol 186, 317–321. 10.1083/jcb.200811067.

39. Borgen, M.A., Wang, D., and Grill, B. (2017). RPM-1 regulates axon termination by affecting growth cone collapse and microtubule stability. Development 144, 4658–4672. 10.1242/dev.154187.

40. Opperman, K.J., and Grill, B. (2014). RPM-1 is localized to distinct subcellular compartments and regulates axon length in GABAergic motor neurons. Neural Dev 9, 10. 10.1186/1749-8104-9-10.

41. Walser, M., Umbricht, C.A., Frohli, E., Nanni, P., and Hajnal, A. (2017). beta-Integrin de-phosphorylation by the Density-Enhanced Phosphatase DEP-1 attenuates EGFR signaling in C. elegans. PLoS genetics 13, e1006592. 10.1371/journal.pgen.1006592.

42. Williams, B.D., and Waterston, R.H. (1994). Genes critical for muscle development and function in Caenorhabditis elegans identified through lethal mutations. J Cell Biol 124, 475–490. 10.1083/jcb.124.4.475.

43. Wang, S., Tang, N.H., Lara-Gonzalez, P., Zhao, Z., Cheerambathur, D.K., Prevo, B., Chisholm, A.D., Desai, A., and Oegema, K. (2017). A toolkit for GFP-mediated tissue-specific protein degradation in C. elegans. Development 144, 2694–2701. 10.1242/dev.150094.

44. Gallegos, M.E., and Bargmann, C.I. (2004). Mechanosensory neurite termination and tiling depend on SAX-2 and the SAX-1 kinase. Neuron 44, 239–249. 10.1016/j.neuron.2004.09.021.

45. Rogalski, T.M., Mullen, G.P., Gilbert, M.M., Williams, B.D., and Moerman, D.G. (2000). The UNC-112 gene in Caenorhabditis elegans encodes a novel component of cell-matrix adhesion structures required for integrin localization in the muscle cell membrane. J Cell Biol 150, 253–264.

46. Poinat, P., De Arcangelis, A., Sookhareea, S., Zhu, X., Hedgecock, E.M., Labouesse, M., and Georges-Labouesse, E. (2002). A conserved interaction between beta1 integrin/PAT-3 and Nck-interacting kinase/MIG-15 that mediates commissural axon navigation in C. elegans. Curr Biol 12, 622–631.

47. Oliver, D., Norman, E., Bates, H., Avard, R., Rettler, M., Benard, C.Y., Francis, M.M., and Lemons, M.L. (2019). Integrins Have Cell-Type-Specific Roles in the Development of Motor Neuron Connectivity. J Dev Biol 7. 10.3390/jdb7030017.

48. Lei, W.L., Xing, S.G., Deng, C.Y., Ju, X.C., Jiang, X.Y., and Luo, Z.G. (2012). Laminin/beta1 integrin signal triggers axon formation by promoting microtubule assembly and stabilization. Cell Res 22, 954–972. 10.1038/cr.2012.40.

49. Eysert, F., Coulon, A., Boscher, E., Vreulx, A.C., Flaig, A., Mendes, T., Hughes, S., Grenier-Boley, B., Hanoulle, X., Demiautte, F., et al. (2021). Alzheimer’s genetic risk factor FERMT2 (Kindlin-2) controls axonal growth and synaptic plasticity in an APP-dependent manner. Molecular psychiatry 26, 5592–5607. 10.1038/s41380-020-00926-w.

50. Huang, C., Rajfur, Z., Yousefi, N., Chen, Z., Jacobson, K., and Ginsberg, M.H. (2009). Talin phosphorylation by Cdk5 regulates Smurf1-mediated talin head ubiquitylation and cell migration. Nat Cell Biol 11, 624–630. 10.1038/ncb1868.

51. Crawley, O., Giles, A.C., Desbois, M., Kashyap, S., Birnbaum, R., and Grill, B. (2017). A MIG-15/JNK-1 MAP kinase cascade opposes RPM-1 signaling in synapse formation and learning. PLoS genetics 13, e1007095. 10.1371/journal.pgen.1007095.

52. Bremer, J., Marsden, K.C., Miller, A., and Granato, M. (2019). The ubiquitin ligase PHR promotes directional regrowth of spinal zebrafish axons. Communications biology 2, 195. 10.1038/s42003-019-0434-2.

53. McCarty, J.H., Lacy-Hulbert, A., Charest, A., Bronson, R.T., Crowley, D., Housman, D., Savill, J., Roes, J., and Hynes, R.O. (2005). Selective ablation of alphav integrins in the central nervous system leads to cerebral hemorrhage, seizures, axonal degeneration and premature death. Development 132, 165–176. 10.1242/dev.01551.

54. Nieuwenhuis, B., Haenzi, B., Andrews, M.R., Verhaagen, J., and Fawcett, J.W. (2018). Integrins promote axonal regeneration after injury of the nervous system. Biol Rev Camb Philos Soc 93, 1339–1362. 10.1111/brv.12398.

55. Hisamoto, N., Shimizu, T., Asai, K., Sakai, Y., Pastuhov, S.I., Hanafusa, H., and Matsumoto, K. (2019). C. elegans Tensin Promotes Axon Regeneration by Linking the Met-like SVH-2 and Integrin Signaling Pathways. J Neurosci 39, 5662–5672. 10.1523/JNEUROSCI.2059-18.2019.

56. Bloom, A.J., Miller, B.R., Sanes, J.R., and DiAntonio, A. (2007). The requirement for Phr1 in CNS axon tract formation reveals the corticostriatal boundary as a choice point for cortical axons. Genes Dev 21, 2593–2606. gad.1592107 [pii]10.1101/gad.1592107.

57. Han, S., Kim, S., Bahl, S., Li, L., Burande, C.F., Smith, N., James, M., Beauchamp, R.L., Bhide, P., Diantonio, A., and Ramesh, V. (2012). The E3 ubiquitin ligase, protein associated with Myc (Pam) regulates mammalian/mechanistic target of rapamycin complex 1 (mTORC1) signaling in vivo through N- and C-terminal domains. J Biol Chem. 10.1074/jbc.M112.353987.

58. Mirdita, M., Schutze, K., Moriwaki, Y., Heo, L., Ovchinnikov, S., and Steinegger, M. (2022). ColabFold: making protein folding accessible to all. Nat Methods 19, 679–682. 10.1038/s41592-022-01488-1.

59. Szklarczyk, D., Gable, A.L., Lyon, D., Junge, A., Wyder, S., Huerta-Cepas, J., Simonovic, M., Doncheva, N.T., Morris, J.H., Bork, P., et al. (2019). STRING v11: protein-protein association networks with increased coverage, supporting functional discovery in genome-wide experimental datasets. Nucleic acids research 47, D607–D613. 10.1093/nar/gky1131.

