## Supplementary figures and images for "Axon development is regulated at genetic and proteomic interfaces between the integrin adhesome and the RPM-1 ubiquitin ligase signaling hub"

### Supplementary Figure 1

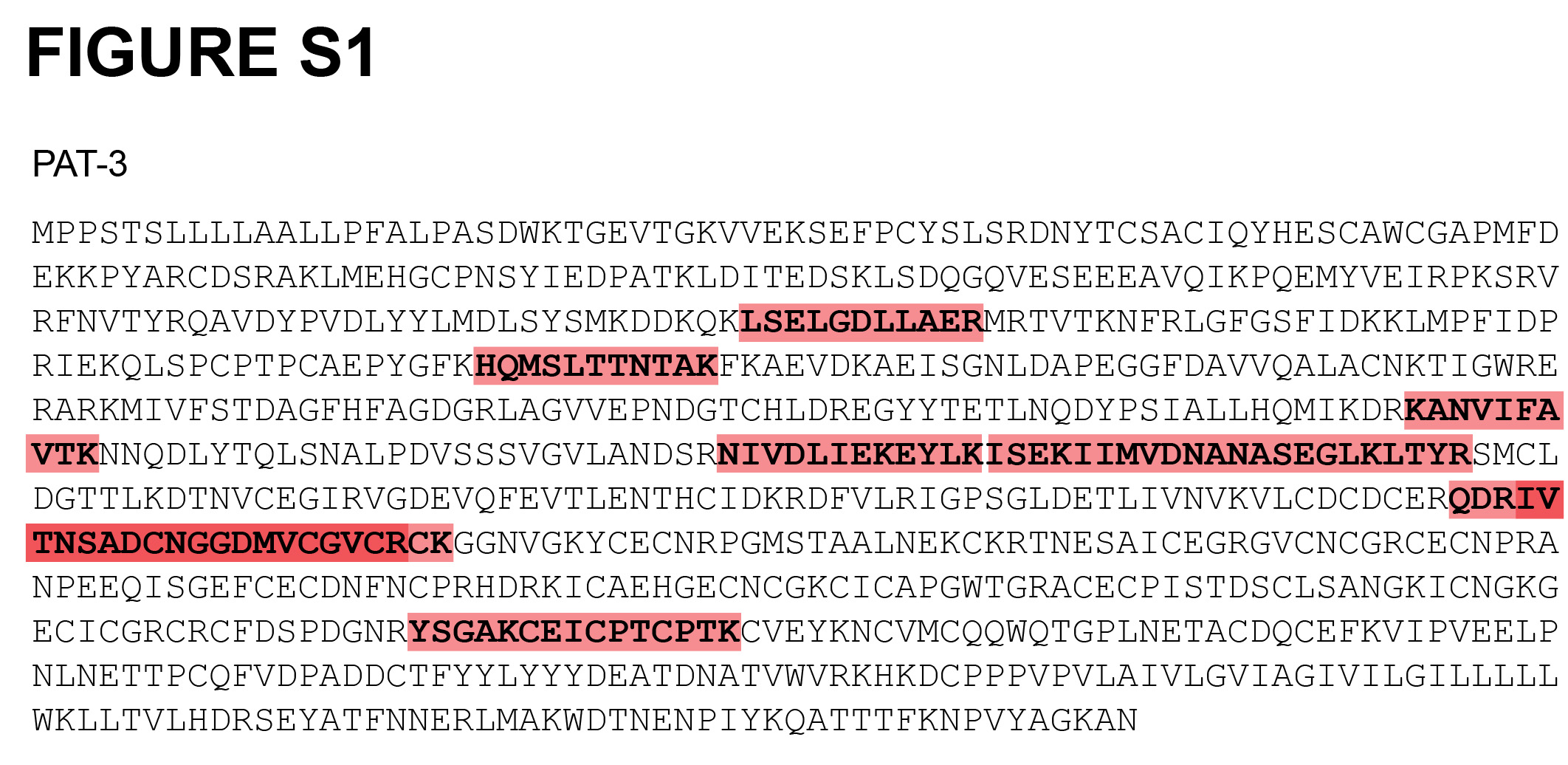

### Supplementary Figure 2

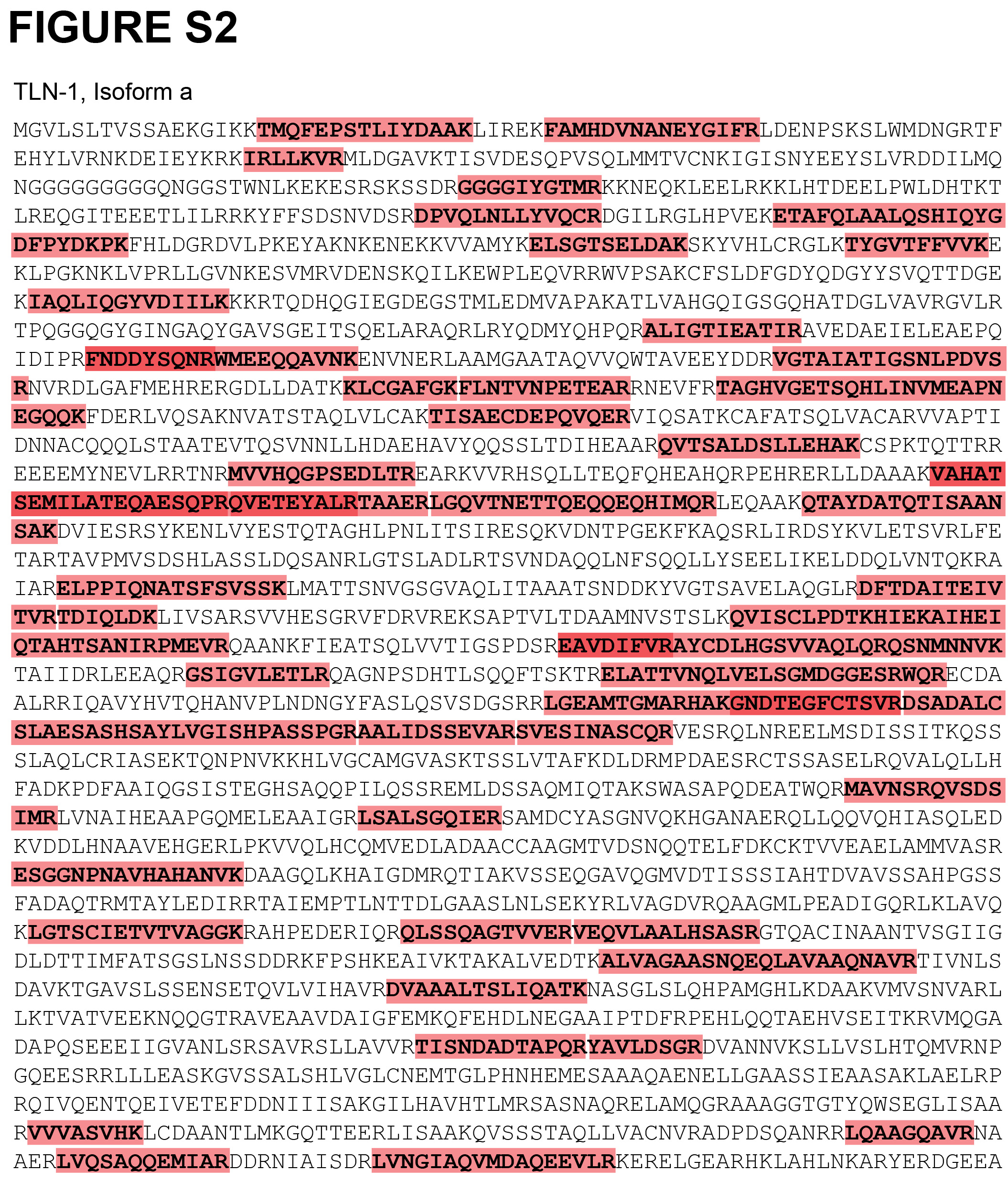

### Supplementary Figure 3

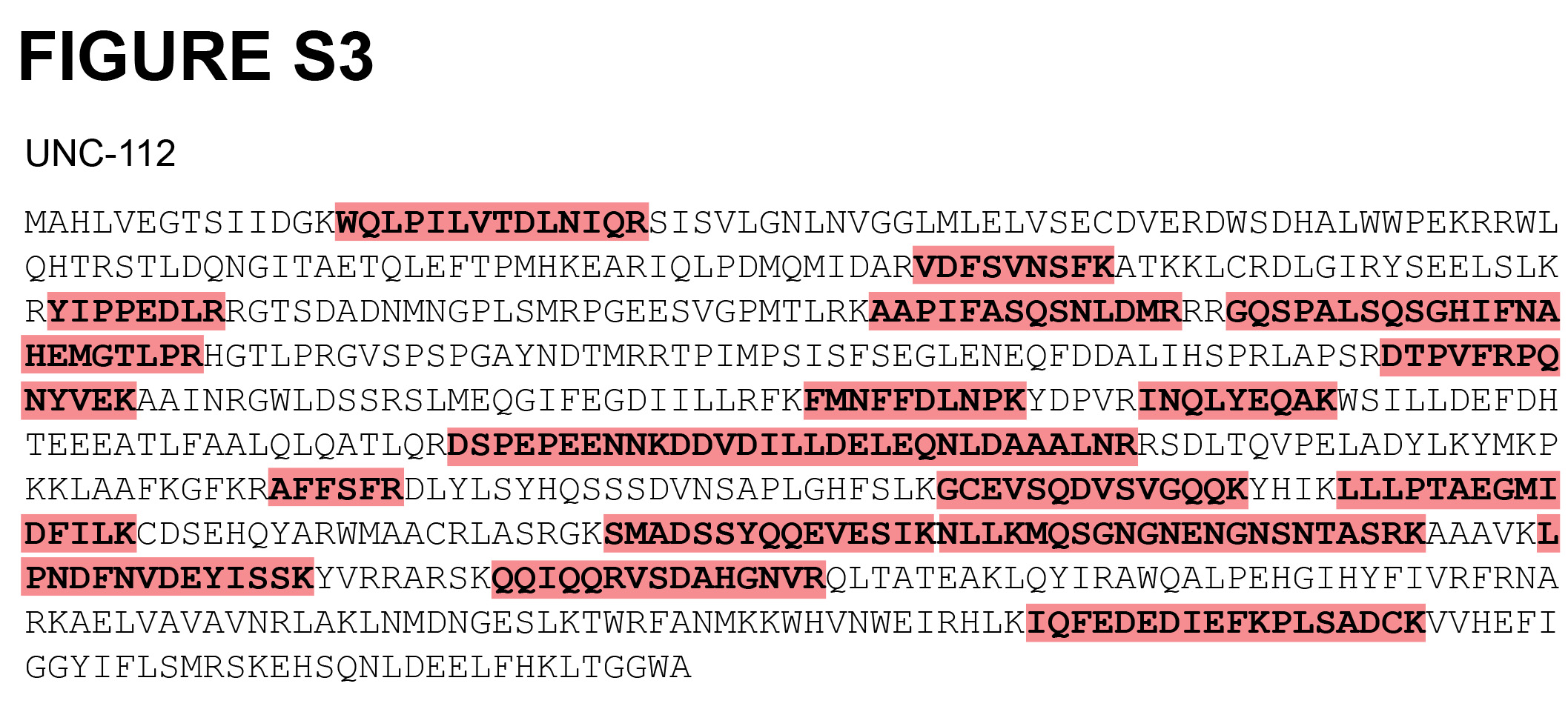

### Supplementary Figure 4

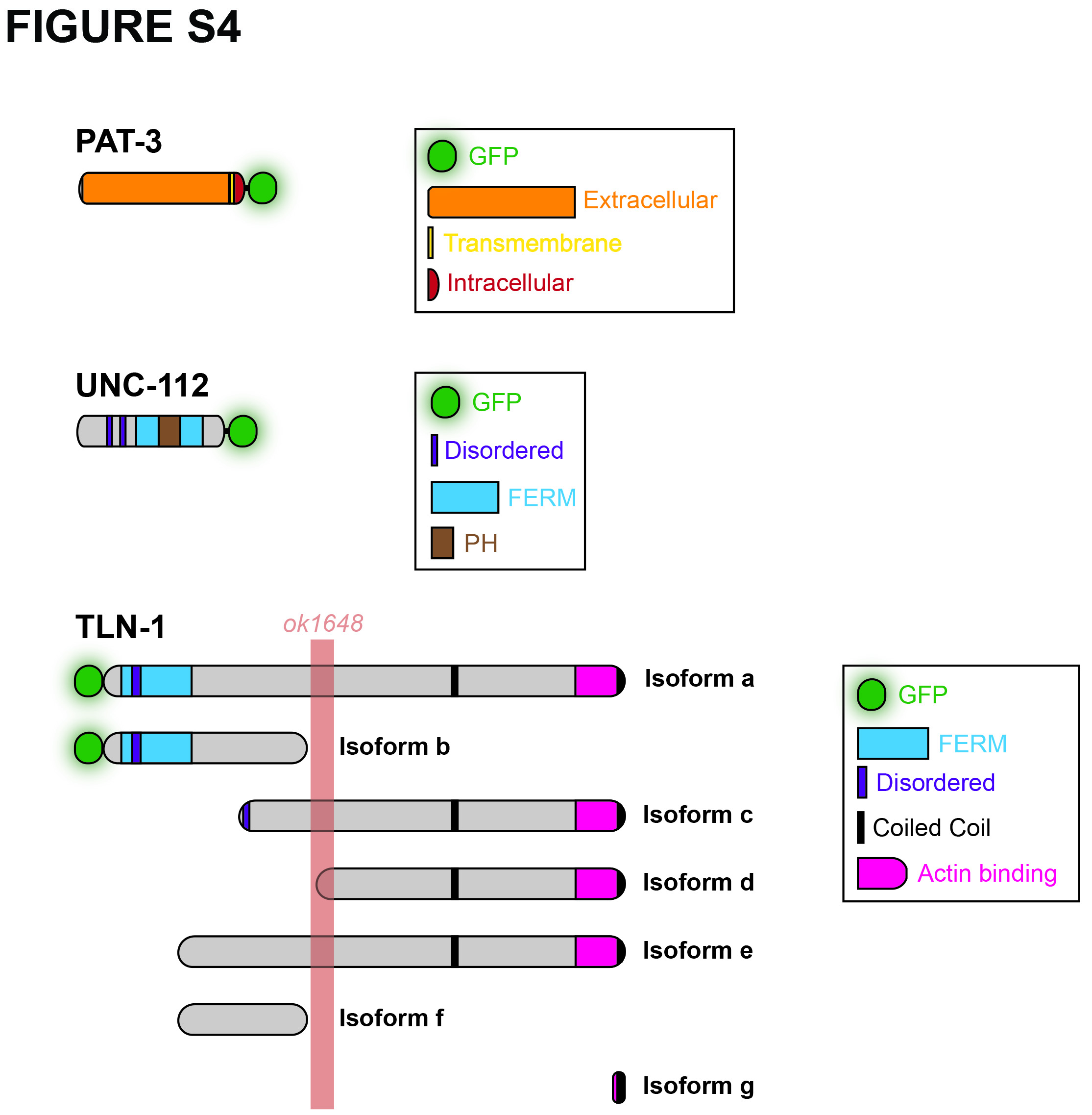

### Supplementary Figure 5

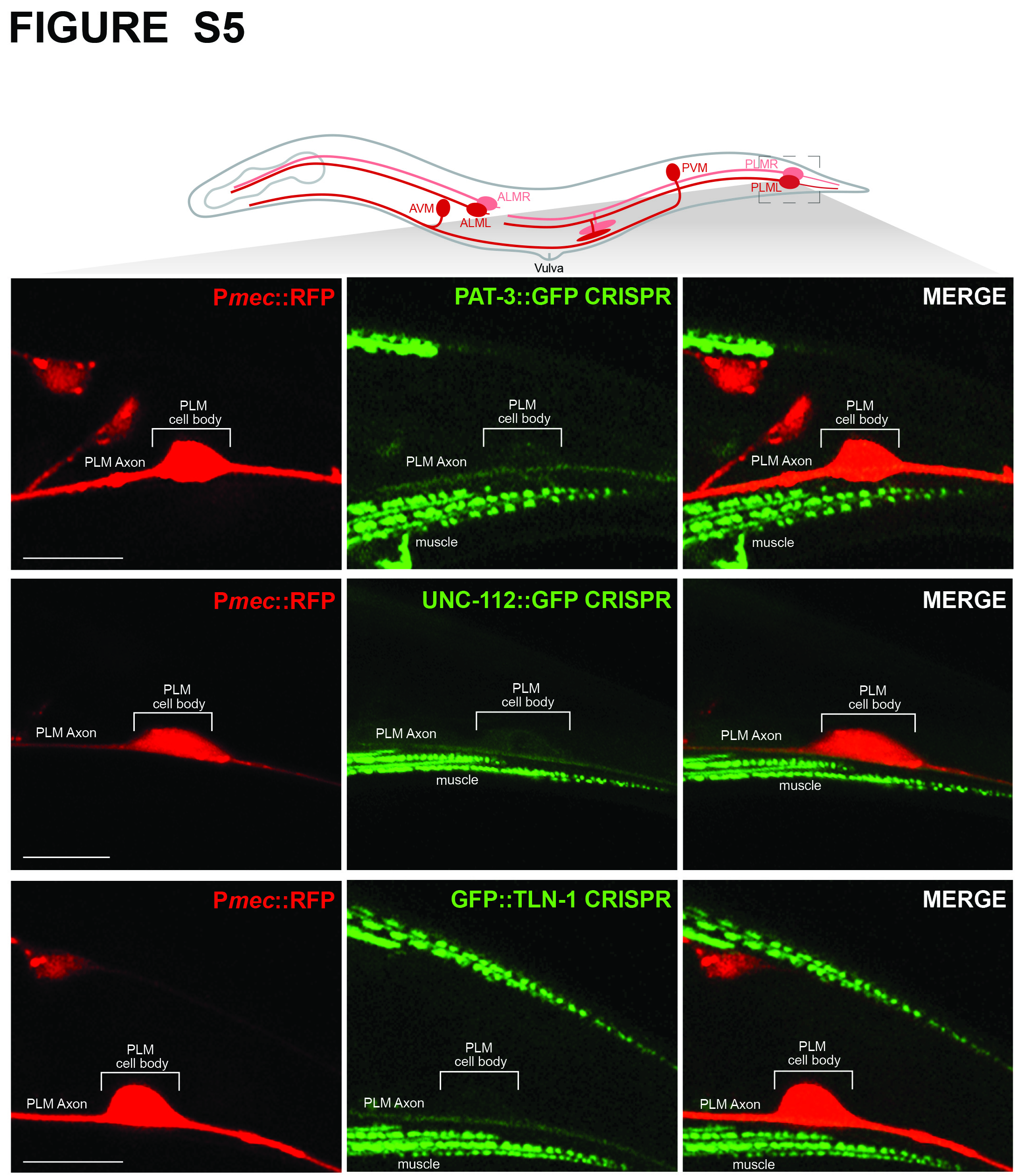
