## Supplementary Table 4 for "Axon development is regulated at genetic and proteomic interfaces between the integrin adhesome and the RPM-1 ubiquitin ligase signaling hub"

**Table S4: Transgenic and CRISPR Strains**

| Figure | Strain | Genotype |
|---|---|---|
| Figure 1, 2 | XMN829 | *muIs32* II; *rpm-1*(*ju44*)<br>*bggIs9* [$P_{rpm\text{-}1}$::GS::RPM-1; $P_{myo\text{-}2}$::mCherry; pha-1(+)] V |
| Figure 1 | XMN830 | *muIs32 bggIs19* [$P_{rpm\text{-}1}$::GS::RPM-1 LD;<br>$P_{myo\text{-}2}$::mCherry; pha-1(+)] II; *rpm-1*(*ju44*) V |
| Figure 1, 2 | XMN831 | *muIs32* II; *rpm-1*(*ju44*) V;<br>*bggIs23* [$P_{rpm\text{-}1}$::GS::GFP; $P_{myo\text{-}2}$:: mCherry, pha-1(+)] |
| Figure 3, 4 | XMN1242 | *pat-3*(*bgg86* [PAT-3::GFP CRISPR]);<br>*jsIs973* [$P_{mec\text{-}7}$::mRFP; unc-119 (+)] III |
| Figure 3, 4, 6 | XMN1231 | *tln-1*(*zh117* [GFP::TLN-1 CRISPR]);<br>*jsIs973* [$P_{mec\text{-}7}$::RFP, unc-119 (+)] III |
| Figure 3, 4 | XMN1168 | *unc-112*(*bgg68* [UNC-112::GFP CRISPR]) V;<br>*jsIs973* [$P_{mec\text{-}7}$::RFP, unc-119(+)] III |
| Figure 4, 5 | XMN913 | *jsIs973*[$P_{mec\text{-}7}$::mRFP] |
| Figure 4 | XMN1167 | *itSi953* [$P_{mec\text{-}18}$::*mecDEG*, unc-119(+)] II;<br>*jsIs973* [$P_{mec\text{-}7}$::RFP, unc-119(+)] III |
| Figure 4, 5 | XMN1243 | *itSi953* [$P_{mec\text{-}18}$::*mecDEG*, unc-119(+)] II;<br>*pat-3*(*bgg86* [PAT-3::GFP])<br>*jsIs973* [$P_{mec\text{-}7}$::RFP, unc-119(+)] III |
| Figure 4, 5, 6 | XMN1230 | *tln-1*(*zh117* [GFP::TLN-1 CRISPR]);<br>*itSi953* [$P_{mec\text{-}18}$::*mecDEG*, unc-119(+)] II;<br>*jsIs973* [$P_{mec\text{-}7}$::RFP, unc-119(+)] III |
| Figure 4, 5 | XMN1359 | *unc-112*(*bgg68* [UNC-112::GFP CRISPR]) V;<br>*jsIs973* [$P_{mec\text{-}7}$::RFP, unc-119(+)] III;<br>*itSi953* [$P_{mec\text{-}18}$::*mecDEG*, unc-119(+)] II |
| Figure 4 | XMN1229 | *tln-1*(*ok1648*); *jsIs973* [$P_{mec\text{-}7}$::RFP, unc-119(+)] III |
| Figure 5 | XMN1152 | *rpm-1(ju44)* V; *jsIs973* [$P_{mec\text{-}7}$::RFP, unc-119(+)] III |
| Figure 5, 6 | XMN1272 | *itSi953* [$P_{mec\text{-}18}$::*mecDEG*, unc-119(+)] II;<br>*jsIs973* [$P_{mec\text{-}7}$::RFP, unc-119(+)] III; *rpm-1(ju44)* V |
| Figure 5 | XMN1366 | *rpm-1(ju44)* V; *itSi953*[$P_{mec\text{-}18}$::*mecDEG*, unc-119(+)] II;<br>*pat-3*(*bgg86* [PAT-3::GFP]);<br>*jsIs973* [$P_{mec\text{-}7}$::RFP, unc-119(+)] III |
| Figure 5, 6 | XMN1305 | *rpm-1(ju44)*; *tln-1*(*zh117* [GFP::TLN-1 CRISPR];<br>*itSi953* [$P_{mec\text{-}18}$::*mecDEG*, unc-119(+)] II;<br>*jsIs973* [$P_{mec\text{-}7}$::RFP, unc-119(+)] III |
| Figure 5 | XMN1258 | *itSi953* [$P_{mec\text{-}18}$::*mecDEG*, unc-119(+)] II;<br>*jsIs973* [$P_{mec\text{-}7}$::RFP, unc-119(+)] III; *rpm-1 (ju44)* V;<br>*unc-112*(*bgg68* [UNC-112::GFP]) V |
| Figure 4 | XMN1412 | *tln-1*(*zh117* [GFP::TLN-1 CRISPR]); *itSi953*<br>[$P_{mec\text{-}18}$::*mecDEG*, unc-119(+)] II; *jsIs973* [$P_{mec\text{-}7}$::RFP,<br>unc-119(+)] III; bggEx172 ($P_{rgef\text{-}1}$::FLAG::TLN-1) |

*mecDEG* = ltSi953 [mec-18p::vhhGFP4::zif-1::operon-linker::mKate::tbb-2 3'UTR + Cbr-unc-119(+)] II
