## Supplementary Table 5 for "Axon development is regulated at genetic and proteomic interfaces between the integrin adhesome and the RPM-1 ubiquitin ligase signaling hub"

### Table S5: Primers for genotyping and cloning

| Gene | Allele/ Construct | Primer Sequence |
|---|---|---|
| *mecDEG* Transgene | *itSi953* | ttTi5605 fwd: 5' TTTCTCAGTTGTGATACGGTTTTT 3'<br>Int: 5' aggaacagaataacagatgatgagc 3'<br>ttTi5605 rev: 5' CGCTACTTACCGGAAACCAA 3' |
| *tln-1* | *ok1648* | ok1648 fwd: 5' gagccaaatgacgagtaggg 3'<br>wt rev: 5' AGAGTCGAACGGATGTTTCG 3'<br>ok1648 rev: 5' tctcatggatcgctttttcg 3' |
| *tln-1* | *zh117* | zh117 fwd: 5' TCCAGATAAACCGCAAATCC 3'<br>zh117 Int: 5' GGCAGACAAACAAAAGAATGG 3'<br>zh117 rev: 5' ATCTGGGCGTGGTTTGTTAG 3' |
| *pat-3* | *bgg86* | bgg86 fwd: 5' TTATGGCCAAATGGGATACG 3'<br>bgg86 rev: 5' TGTGAGTTGTTGGTCGGTGT 3'<br>bgg86 Int: 5' GGCAGACAAACAAAAGAATGG 3' |
| *unc-112* | *bgg68* | wt fwd: 5' CGAAGCAAAGAACATTCTCAA 3'<br>bgg68 fwd: 5' GGCAGACAAACAAAAGAATGG 3'<br>wt rev: 5' TTGTACCTACGTTTGCCTAC 3' |
| *rpm-1* | *ju44* | *ju44* fwd: 5' CGTGTATGACCTGTAAACGAGAAG 3'<br>*ju44* rev: 5' GACATGTTGGAAGAAGATGTTTTG 3'<br>Digest with AccI |
