## Supplementary Table 6 for "Axon development is regulated at genetic and proteomic interfaces between the integrin adhesome and the RPM-1 ubiquitin ligase signaling hub"

### Table S6: Injection Conditions

| Figure | Transgene | Injected Strain | Injection Mix |
|---|---|---|---|
| **Figures 3, 4, 5** | PAT-3::GFP CRISPR (*bgg86*) | **N2** | 5µL tracrRNA (4µg/µL) |
| | | | 0.4µL dpy-10 crRNA (8µg/µL) |
| | | | 0.55µL dpy-10 repair ssODN (500µg/µL) |
| | | | 1µL PAT-3::GFP crRNA (8ug/µL) |
| | | | 6.8µL PAT-3::GFP PCR Template (500ng/µL) |
| | | | 0.5µL KCl (1M) |
| | | | 0.75Ml Hepes pH 7.4 (200mM) |
| | | | 4.6µL Nuclease-Free $H_20$ |
| | | | 5µL Cas9 (10µg/µL) |
| **Figures 3, 4, 5** | UNC-112::GFP CRISPR (*bgg68*) | **N2** | 5µL tracrRNA (4µg/µL) |
| | | | 0.4µL dpy-10 crRNA (8µg/µL) |
| | | | 0.55µL dpy-10 repair ssODN (500µg/µL) |
| | | | 1µL UNC-112::GFP crRNA (8ug/µL) |
| | | | 6.8µL UNC-112::GFP PCR Template (500ng/µL) |
| | | | 0.5µL KCl (1M) |
| | | | 0.75Ml Hepes pH 7.4 (200mM) |
| | | | 4.6µL Nuclease-Free $H_20$ |
| | | | 5µL Cas9 (10µg/µL) |
| **Figure 4** | TLN-1 Rescue | *tln-1*(*zh117*[GFP::TLN-1]); *itSi953*[$P_{mec-18}$ *mecDEG*, *unc-119*(+)] II; *jsIs973*[$P_{mec-7}$::RFP, *unc-119*(+)] III | 65ng/µL pBluescript (pBG-49) |
| | | | 10ng/µL Prps-27::NeoR (pBG-264) |
| | | | 25ng/µL Prgef-1::FLAG::TLN-1 (pBG-GY1100) |
