## Supplementary Table 7 for "Axon development is regulated at genetic and proteomic interfaces between the integrin adhesome and the RPM-1 ubiquitin ligase signaling hub"

### Table S7: CRISPR Targeting Sequences and Repair Templates

| Gene | crRNA | crRNA Target Sequence | Repair Template |
|---|---|---|---|
| *unc-112* | UNC-112::GFP | CTTTTCCACAAACTTACAGG | GCAAAGAACATTCTCAAAATCTTGATGAAGAACTTTT CCATTAAGCTCACTGG***AGG***ATGG GCTGGAGGAGGAGGATCCGGAGGAGGAGGATCCGGA GGAGGAGGATCC --- GFP --- TAGATATTTAAATTTCTATAATCTTTTGCAAACCA |
| *pat-3* | PAT-3::GFP | ACTAAATAGTTTTTATCCTT | Hybrid dsDNA made of a PCR with no gene homology and PCR with a long homology repair template<br><br>No gene homology repair template<br>GGAGGAGGAGGATCTGGTGGTGGAG ---GFP--- TAA<br><br>long homology repair template<br>AAGTGACACGGAACAAAAAACATATAAATTTATCAAA TTATCATTTTTCAGAACGAGAACCCAATCTACAAACA GGCCACGACAACATTTAAAAATCCAGTATACGCTGGA AAAG***CCA***ACGGAGGAGGAGGATCTGGTGGTGGAGGA TCTGGTGGAGGTGGATCA ---GFP--- TAAATAGTTTTTATCCTTATATTTTAATAATTTTCCCAA ATTTTCTAATATGAAAGCTCAATTTCTCCATCCAAACA ACTCGAAACGAGTATTTAGCGATAAATTGTCACATTCT TCTTTGTTTTATTCAAAAAATCT |

Legend: underline (crRNA targeting sequence), Italic bold (Pam sequence), Red (Insertion), Blue (silent mutations in repair to prevent Cas9 re-cutting), Orange (linker)
